# Temporal dynamics of RNA metabolic enzyme interactome lay the foundation for riboregulation mediated circadian metabolism

**DOI:** 10.1101/2025.07.31.667909

**Authors:** Dinesh Balasaheb Jadhav, Sougata Roy

## Abstract

The temporal regulation of the RNA-binding to metabolic enzymes offers a compelling framework to address the open question of whether circadian control of metabolism can be mediated through riboregulation. To advance this understanding, we investigated the time-of-the-day dependent dynamics of RNA-enzymes interactome using a time-resolved RIC approach. 70% of the captured RBPs exhibit differential RNA-binding between subjective day and night, with 27% associated to metabolic pathways, suggesting temporal riboregulation of metabolic enzymes can be widespread. Intriguingly, 63 metabolic enzymes displayed circadian RNA-binding uncoupled from their protein abundance dynamics. This observation suggests that the temporal regulation of their RNA-binding “moonlighting” functions is likely mediated by alternative mechanisms, such as post-translational modifications. Our study presents a catalog of metabolic enzymes localized in mitochondria and chloroplast that exhibit dynamic RNA binding across circadian time, providing a valuable testbed to explore the mechanistic role of riboregulation in driving daily metabolic rhythms across species.

## Introduction

Daily metabolic rhythms are conserved across a wide range of species and are driven by a self-sustaining, cell-based circadian clock. When synchronized with environmental cues, the circadian clock enhances organismal fitness by anticipating daily changes and optimizing the allocation and utilization of cellular resources^1–4^. While the core components of the circadian clock are well characterized across taxa^5–10^, the mechanisms by which the clock integrates and orchestrates metabolism across the daily cycle remain a major open question. To elucidate the mechanistic underpinnings of circadian regulation of metabolism, a conventional approach has been to investigate the temporal dynamics of metabolic enzymes at the mRNA and protein levels, as these are considered direct proxies for the output of their corresponding metabolic pathways. Consequently, circadian transcriptomics^11–16^, proteomics^17–21^, and analyses of circadian posttranslational modifications^22–26^ have been the most widely used strategies to uncover clock-mediated metabolic regulation. Apart from providing excellent insights to the clock regulation of metabolism, this course of investigation led us to realize that activity of enzymes is not always proportional to its abundance. There may be several reasons behind it, one that strike us and encouraged us to investigate it further was the process of riboregulation, a process where RNAs regulate the function of their cognate proteins. RNA-binding proteins (RBPs) were studied for their ability to modulate the function of the RNAs they associate with^27–29^. However, recent breakthroughs in RNA-interactome technologies have revealed a wide array of metabolic enzymes with RNA-binding “moonlighting” functions, an area that has garnered significant attention^30–32^. A growing consensus has highlighted the moonlighting roles of RBPs across diverse organisms and in different physiological contexts, including cellular energetics^33–36^, DNA biosynthesis^37^, amino acid metabolism^38^, lipid metabolism^39^, folate metabolism^40^, cell division^41^, viral infection^42^, cardiovascular biology^43^, autophagy^44^ and stem cell differentiation^34^. Given its critical role in regulating cellular physiology and metabolism^31,32^, we anticipate that riboregulation will play a major role in the circadian regulation of cellular metabolism.

Carbohydrate metabolism, for instance, is under circadian control in mammals^45,46^, plants^1,47^, and algae^48,49^ at both systemic and cellular levels. In mammals, disruption of the suprachiasmatic nucleus impairs glucose homeostasis^50,51^, and similar temporal regulation is observed in glycogen metabolism^52–54^ and gluconeogenesis^55,56^. In the green lineage, carbohydrate metabolism is tightly integrated with photosynthesis, a process regulated by the circadian clock^1,57^, which remains conserved across diverse taxa, including angiosperms, bryophytes^58^, liverworts^59^, and algae^60–63^. Beyond carbohydrate metabolism, the circadian clock also regulates a broad range of fundamental metabolic processes, including lipid, protein, amino acid, redox, and cellular energetics pathway, highlighting its central role in maintaining metabolic homeostasis across diverse taxa^49,64–66^. Notably, many of the enzymes involved in the fundamental metabolic processes mentioned above are not only highly conserved across kingdoms, underscoring their central role in core metabolic pathways shared among species but also exhibit RNA-binding moonlighting function. What particularly intrigued us was the ability of the protein bound RNAs to regulate the enzymatic activity of these metabolic enzymes^34,35,38,67^. For example, recent studies have shown that RNA binding to the glycolytic enzyme enolase 1 can modulate its catalytic function and alter the cellular glycolysis output, thereby influencing embryonic stem cell differentiation in humans^34^. Similarly, malate dehydrogenase 2 (MDH2), a key enzyme in the TCA cycle, binds RNA in a manner that may compete with its cofactor NAD⁺^35^, a mechanism consistent with previous findings in other metabolic enzymes^68^. In *Chlamydomonas reinhardtii*, the dihydrolipoamide acetyltransferase subunit (DLA2), a component of the pyruvate dehydrogenase complex localized in chloroplast was found to shift from its canonical enzymatic role to an RNA-binding function in response to a post-translational modification^67^. If RNA binding to the metabolic enzymes can modulate its catalytic function, it is plausible that its activity could be temporally regulated by the underlying circadian dynamicity of RNA-binding.

We identified a gap in the current understanding of circadian riboregulation, highlighting the need for a comprehensive catalogue of proteins and enzymes exhibiting circadian RNA-binding activity as a foundational step. The RNA–RBP interactome across the 24-hour circadian cycle remains largely unexplored. As a first step toward addressing this gap, we were interested in how dynamic is the RNA–RBP interactome over a daily cycle? Using circadian RNA-interactome capture approach we investigated whether metabolic enzymes exhibit time-of-day dependent changes in their moonlighting RNA-binding functions. We hypothesize that the temporal dynamics of RNA-metabolic enzyme interaction may contribute to the circadian regulation of their enzymatic activity. To test this, we employed a simple and experimentally tractable unicellular model system of *Chlamydomonas reinhardtii*, which is well established for studying clock-controlled mechanisms due to its diverse and readily measurable circadian physiology^69^. Additionally, *C. reinhardtii* is a unicellular alga that exhibits features reminiscent of both plant and animal systems^70^. Due to its close evolutionary relationship to the last common ancestor of the plant and animal lineages^71^, elucidating the molecular architecture and riboregulatory dynamics of the *C. reinhardtii* circadian system may provide key insights into the evolutionary origins and diversification of eukaryotic circadian riboregulation. Moreover, given that its circadian clock components are characterized^69,72–74^ and both diurnal and circadian proteomic^13,17^ datasets are available further supports its utility for such investigations.

Using *C. reinhardtii*, we assessed the dynamic nature of RNA–RBP interactions under circadian conditions. Using a previously published, unbiased RNA-interactome capture approach, we identified both RBPs and their associated protein-bound RNAs (PBRs), followed by high-throughput quantitative transcriptomic and proteomic analyses. Specifically, we identified 518 RBPs in *Chlamydomonas reinhardtii*, of which approximately 27% are classified as metabolic enzymes. Notably, we observed strong evolutionary conservation of RNA-binding metabolic enzymes from *C. reinhardtii* to humans. Nearly 70% of these RBPs exhibited circadian time-dependent alterations in their RNA-binding activity, with about one-quarter being metabolic enzymes. Many temporally active RBPs originate from mitochondria and chloroplasts, key hubs of cellular energetics and cellular metabolism, which are themselves regulated by the circadian clock. Given their pivotal roles, they are likely critical in coordinating temporal patterns of metabolism. We asked whether the RNA-binding activity of RBPs is proportional to their abundance. This prompted us to investigate whether the dynamic RNA-binding behaviour observed in metabolic enzymes is directly correlated with their protein levels. To assess this, we focused on identifying “circadian dynamic binders”, proteins whose RNA-binding activity fluctuates with a circadian pattern independently of their abundance profiles. We examined 396 proteins for which both abundance and RNA-binding intensity data were retrievable. Of these, nearly 65% (255) were classified as circadian dynamic binders, and one-third of those were metabolic enzymes. Our catalogue of dynamic binders includes key proteins involved in core metabolic pathways such as photosynthesis, glycolysis/gluconeogenesis, TCA cycle, amino acid metabolism, carbon metabolism, fatty acid metabolism, and reactive oxygen species (ROS) metabolism. Based on their RNA-binding profiles, we further categorized these metabolic enzymes as predominantly day- or night-active. We have uncovered a wide array of metabolic enzymes in which temporal dynamics of RNA-binding may play a role in the circadian regulation of metabolism. Our results not only provide the first comprehensive RBP repertoire for the phytoplankton clade but also offer valuable insights toward our overarching goal of understanding circadian riboregulation.

## Results

### RIC successfully apprehends the RBPome of *Chlamydomonas*

The schematics (Figure 1A) outlines the overall workflow of our approach. As this is the first report of RNA interactome capture (RIC) from phytoplankton, we first optimized UV-induced RIC in *C. reinhardtii*. After crosslinking the cells with varying doses of UV, the RNA-bound proteome was extracted, resolved on SDS-PAGE, and visualized by silver staining (Figure 1B, top panel). Alternatively, following SDS-PAGE, western blotting was performed using an antibody against a known RNA-binding protein (RBP40) (Figure 1B, bottom panel). Both approaches confirmed that 4.55 J/cm² is the optimal UV dose for effective RNA-RBP crosslinking. We further verified the crosslinking efficiency (CE) using the ratio of free and protein bound RNAs, following procedures described in a previous study^75,76^. We found 4.55 J/cm² effectively crosslinked RNAs to RBPs with a CE of approximately 75% (Figure 1C and supplementary figure 1A), without compromising cell integrity (Supplementary Figure S1B) or affecting the total RNA quantity (Figure 1C). Further quality control of the interactome was performed by immunoblotting and silver staining of the different organic phases (Supplementary figure 1C and D). To verify that UV crosslinking effectively enriches RBPs, we employed two validation strategies that were complementary. First, we used an RBP40 antibody to probe RNA interactome captures from crosslinked (CL) and non-crosslinked (NC) cells using immunoblotting. A signal was detected exclusively in CL samples, demonstrating UV-induced RBP enrichment (Figure 1D). Second, we treated samples with RNase to deplete the RNAs, reasoning that no detectable protein bands should appear in the RBP eluates of RNA-depleted samples. Indeed, silver-stained SDS-PAGE analysis showed almost no detectable proteins in the RNase-treated CL sample lanes, confirming that the enriched proteins were RBPs (Figure 1E). Using the same pool of RIC material, we captured PBRs at subjective dawn and subjective midnight to assess their temporal regulation. For simplicity, only mRNAs were enriched from the PBRs using oligo (dT) beads and subsequently sequenced.

**Figure 1:**
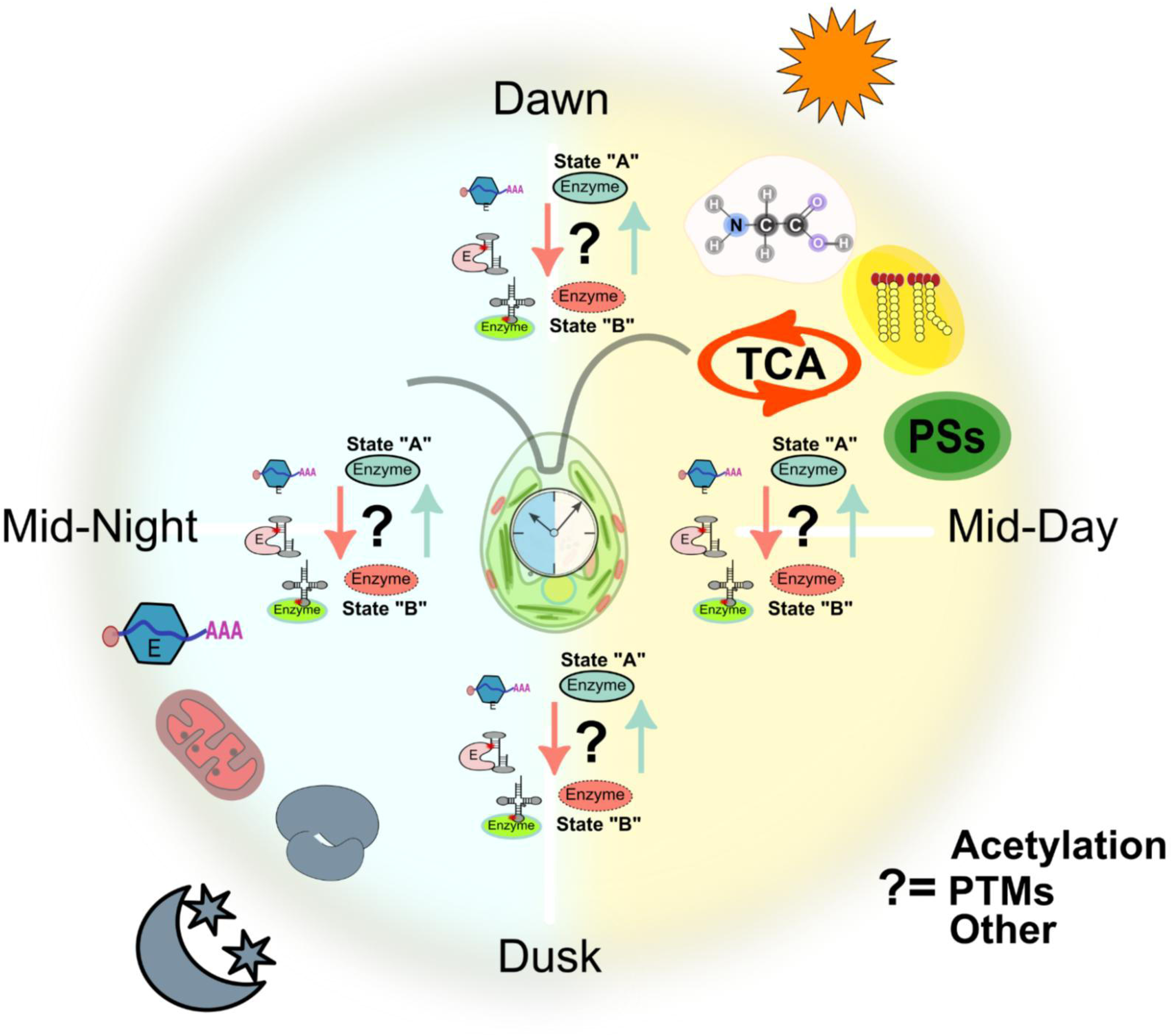
Capturing the circadian RNA interactome from *Chlamydomonas reinhardtii*. **1A)** Schematic representation of circadian RNA interactome capture at 4-time points (LL-48, LL-54, LL-60, and LL-66) and the subsequent downstream processing to quantitative proteomics and transcriptomics. **1B) Top:** A 10% polyacrylamide gel with SDS was silver stained and showing the RBP capture at NC (Non-crosslinked) and crosslinking (CL) at different UV doses indicated at the lop of the respective lanes. The bottom band indicated by a black arrow represents the RNase. **Middle:** Immunoblot showing the recovery of a known RNA binding protein RBP-40 recovery at the same UV doses as mentioned in 1B top panel. **Bottom:** Histogram depicting the relative quantification of the RBP-40 at 5 different UV doses. The Y-axis represents relative intensity. The relative intensity is calculated using the BIORAD ImageLab software. The X-axis represents the the respective UV doses. **1C) Left and right:** Satterplot representing the percentage recovery of the protein bound RNA in CL and NC samples. Two replicates from each time point i.e 8 CL and 8 NC samples were used to test the percentage recovery of protein-bound RNA (PBR) and the total RNA. The red dot represents the mean of the 8 samples and the error bar represents the standard deviation. (***) represent the significant differences and ns stands for not significant. **1D) Top:** Immunoblot illustrating the recovery of RBP-40 at 4 time points in the CL and the NC samples. **Bottom:** Histogram showing the relative intensity of the RBP-40 at 4-time points in the CL and the NC samples.**1E)** 10% PAGE with SDS silver stained gel illustrating the recovery of RBPs from NC and CL samples with and without RNase treatment.Left: Lane 1- NC sample replicate 1, Lane 2- NC sample replicate 2. Lane 3-NC sample with RNase treatment replicate 1, Lane 4-NC sample with RNase treatment replicate 2, Lane 5- Protein marker. Lane 6- CL sample replicate 1, Lane 7 - CL sample replicate 2. Lane 8- CL sample with RNase treatment replicate 1, Lane 9 - CL sample with RNase treatment replicate 2. The replicate 1 and 2 are two experiments with high and low starting cell density, respectively **1F)** Volcano plot depicting the RBPs enriched by comparing Dawn CL vs NC RIC experiment (n=3) using log2 fold difference on X-axis and its significance -log10 (adj_p_value) on Y-axis. A cut off of FDR <0.05 and CL/NC log2 Fold difference of >=1 and <=2 are considered for candidate RBPs (blue coloured), and for RBPs (Coral colored) the cut off was CL/NC FC >=2. Non-significant proteins are colored grey. **1G)** Histogram showing the comparison of the percentage (No of RBPs/Total Proteins) of RBPs identified in *C.reinhardtii* and other species. **1H)** Histogram illustrating the GO enrichment of the RBPs.The Y-axis shows GO-terms. The blue colored bar represents GOBP, green are GOCC and orange are GOMF. Y-axis shows the Fold enrichment. All the terms are BH-FDR <0.05. **1I)** Scatterplot showing the enrichment of the domains. The size represents fold enrichment and the color represents the -log10 adjusted p-value.

*C. reinhardtii* cells were synchronized under 12 h:12 h light-dark cycles for at least five days before being transferred to constant light conditions to free-run for 48 hours. At defined circadian times, the RNA interactome was fixed in vivo using our optimized protocol (Figure 1A), followed by quality validation through immunoblotting (Supplementary Figure 1E). RNA-binding proteins (RBPs) were subsequently isolated from both crosslinked (CL) and non-crosslinked (NC) samples, each in biological triplicates and submitted to the proteomics facility at CSIR Institute of Genomics and Integrative Biology, India for DIA-SWATH-MS analysis. To utilize SWATH-MS for datasets, the primary requirement is to generate a spectral library to serve as a reference for quantification. We identified 2,597 spectra, 1,988 distinct peptides, and 653 proteins, applying a standard 1% FDR at the PSM and peptide level. Using this library, for each circadian timepoints we identified RBPs based on a criterion of CL/NC > 2 with an FDR < 0.05, and classified potential RBPs when 2 > CL/NC > 1, also with an FDR < 0.05 (Figure 1F and Supplementary Figure S1F). In rare cases where no peptides were detected in NC samples but were present across all CL replicates, we manually added these proteins to the RBP list, as the CL/NC ratio would otherwise not yield a valid score. To comprehensively map the RBPome of *C. reinhardtii*, we combined from all four time points (three replicates each for CL and NC samples). In total, we identified 518 RBPs from *C. reinhardtii* (creRBPs), representing the first characterization of an RBPome in a phytoplankton species (Figure 1G). The proportion of creRBPs (RBP/total protein) is approximately 2.7%, comparable to that in *Drosophila melanogaster*, but higher than that reported for the plant *Arabidopsis thaliana* and the worm *Caenorhabditis elegans*. In contrast, the percentage of RBPs in mammals (mouse and human) is nearly twice that observed in *C. reinhardtii* (Figure 1G). To assess whether our methodology introduced bias toward specific types of RBPs, we compared the physicochemical properties such as molecular weight, isoelectric point, and hydrophobicity of the 518 captured RBPs with those of the total proteome. This analysis confirmed that the capture technique is unbiased with respect to these properties (Supplementary Figure S1G). RNA-binding proteins (RBPs) are known to possess extended intrinsically disordered regions (IDRs), which are regions within the protein that lack a stable three-dimensional structure. A comparative analysis of the distribution of disordered regions in creRBPs, and the complete proteome, revealed that creRBPs show a higher proportion of disordered regions (Supplementary Figure S1H). Next, we categorized the 518 creRBPs based on their biological processes, catalytic functions, subcellular localization, and domain architecture. The RNA-binding nature of these proteins is clearly supported by the molecular functions and biological processes associated with the enriched creRBPs (Figure 1H). We identified numerous enzymes involved in metabolic pathways such as photosynthesis, TCA cycle, glycolysis, amino acid and fatty acid metabolism that bind RNAs (Figure 1H), highlighting their moonlighting functions, as observed in previous RIC studies across various species. As expected, known RNA-binding domains were overrepresented among the enriched proteins (Figure 1I). Additionally, our dataset revealed a substantial number of RBPs containing Rossmann folds and P-loop NTPase domains. Moreover, we identified several proteins lacking any known domains, suggesting that many RNA-binding motifs in *C. reinhardtii* remain to be defined.

### Conserved RNA-Binding Moonlighting of Metabolic Enzymes from Humans to Alga

We compared the biological processes, molecular functions, and cellular components of creRBPs to the available datasets from mammals (human and mouse), plant (*Arabidopsis thaliana*), worm (*Caenorhabditis elegans*), insect (*Drosophila melanogaster*) and fungi (*Saccharomyces cerevisiae*). The RBP details of the other species were obtained from RBPbase (https://apps.embl.de/rbpbase/). creRBPs showed species-wide conservation of key RBP related processes and functions (Figure 2A). We also found some unique functions associated with creRBPs such as photosynthetic electron transport (PET) in photosystem I (PSI), dicarboxylic, fatty acid and acetyl coA biosynthesis pathway, thereby expanding the moonlighting function repertoire of RBPs. We also compared the canonical and non-canonical domains of *C. reinhardtii* RBPs with that of other species. To perform this analysis, we uploaded the creRBPs along with RBPs from other species to the STRING database, which directly provided enriched protein domains with a significance threshold of *P* < 0.05. The enriched domain from algae, plant and human showed many common and unique *C. reinhardtii* specific domains (Figure 2B). creRBPs seem to have extensive affiliation with a wide range of cellular processes. We compared the percentage of proteins involved in particular biological pathways that have RNA-binding function and termed it as “coverage”. Specifically, we showed the common RNA binding functions that are conserved across species (Figure 2C) and the insights into mitochondria (Figure 2D) and chloroplast (Figure 2E) associated biological processes. For most of the RNA-binding related functions and the mitochondria related processes creRBPs coverage is similar to that of plants except for ribosome and rRNA binding functions, where creRBPs coverage is higher than that of plants. In chloroplast associated functions except for the photosynthetic electron transport chain, all other processes have almost similar coverage between plants and *C. reinhardtii*. In *C. reinhardtii,* a higher percentage of proteins affiliated with the Photo-ETC show RNA-binding.

**Figure 2:**
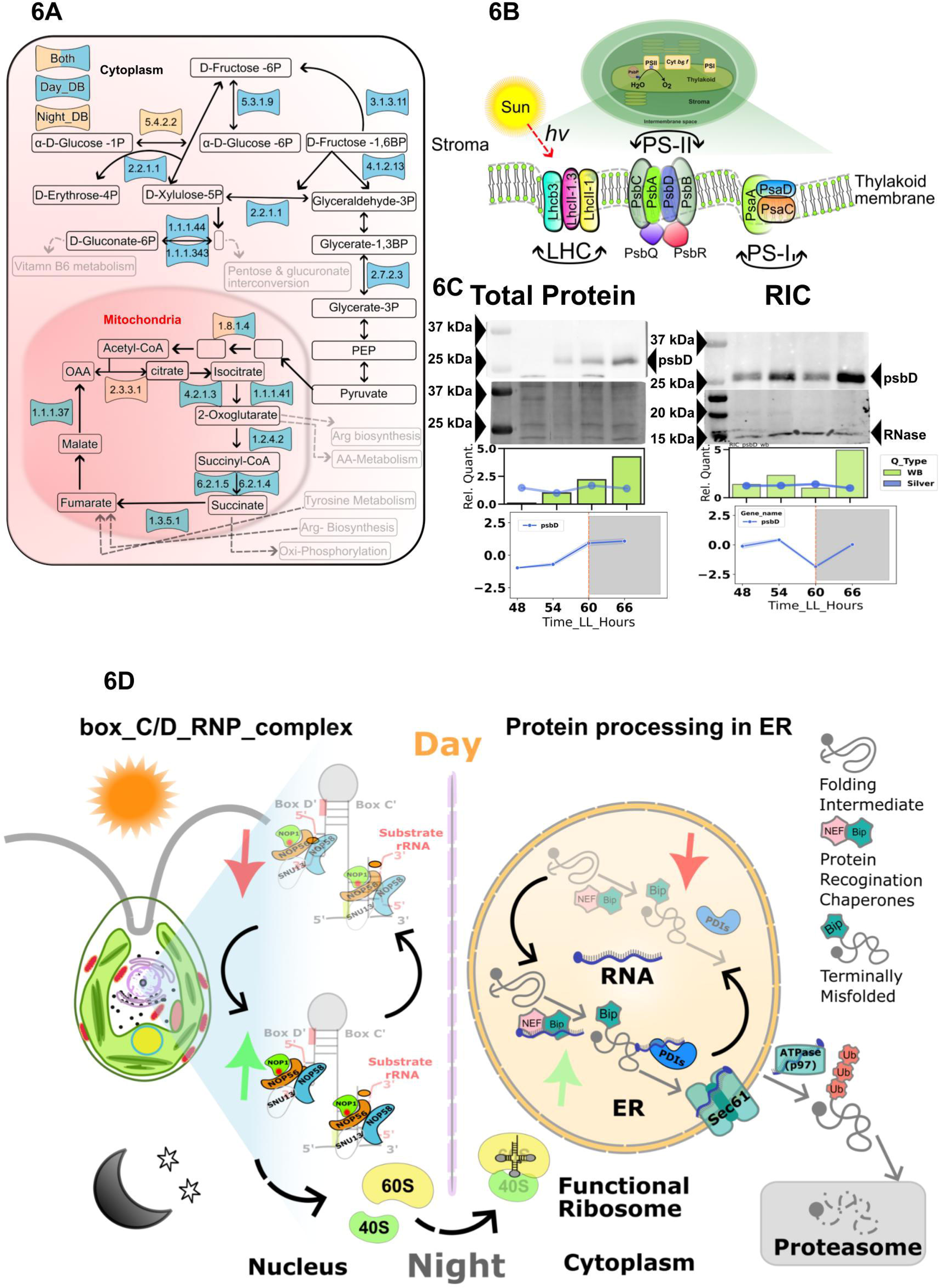
Conservation of RBPs across the species. **2A)** Scatterplot representing the comparison of the GO-enriched terms across the species. The RBPs identified in different species was downloaded from (RBPbase v0.22 alpha).Using the Panther gene ontology, we performed the GO enrichment analysis for all the species. After getting the enrichment, we mapped the unique and the common terms present in these species. Finally, we created this plot in R with the BH corrected FDR and Fold enrichment. The X-axis represents the species and the Y-axis shows the GO-Terms. The size and colour of the filled circles represents fold enrichment and FDR values respectively. All the terms have BH-FDR <0.05.**2B)** Scatterplot illustrating the Fold enrichment of the domains across the algae, plants, and humans. We calculated fold enrichment using stringDB. The green-filled square represents humans, the yellow cross represents the algae and the blue squares represent the plants. **2C) Top:** Scatter plot showing the percentage coverage of individual GO terms from different species, where % coverage refers to the proportion of proteins within each GO-term pathway that were identified as RBPs. Organisms are represented by distinct shapes and colors, as indicated in the figure legend. **2D)** Species-wide comparison of proportion of proteins associated with the GOCC:Mitochondria that were identified as RBPs, which is expressed as % coverage. **2E)** Comparison of proportion of proteins associated with the GOCC:Chloroplast that were identified as RBPs, which is expressed as % coverage in plants and algae. **2F)** Histogram representing the percentage of the RBPs associated with the metabolic pathways in different organisms. **2G)** Scatter plot illustrating the species-wide distribution of metabolic enzymes from the Glycolysis, **2H)** TCA cycle **2I)** oxidative phosphorylation, and **2J)** fatty acid metabolism pathways that exhibit RNA-binding activity. RBPs from all the organism were downloaded from the RBPbase. These RBPs were then searched in KEGG mapper. The color scheme is consistent with Figure 2D.

RNA-binding as a moonlighting function of metabolic enzymes is evident across higher eukaryotes. The evolutionary conservation of these moonlighting enzymes from humans to unicellular algae would further ensure its primordial origin. The preservation of moonlighting function suggests its importance and significance across distant species. In this context, we chose *C. reinhardtii*, which is unique because of its association with the common ancestor of both plants and animals. To reveal the extent of the conservation, we compared the metabolic enzymes across species that demonstrate moonlighting RNA-binding function using the KEGG mapper. Overall, we found almost 27% (140/518) of the captured creRBPs are affiliated with metabolic pathways, whereas in humans and plants, this affiliation is only 11% and 18%, respectively (Figure 2F and Supplementary Figure 2A). This suggests the RNA binding as a moonlighting function of metabolic enzymes is widespread in the green algae *C. reinhardtii*, which indicates its importance as a model to find novel riboregulatory functions.

Our analysis shows an interesting pattern: the moonlighting function of enzymes involved in glycolysis (Figure 2G) and the TCA cycle (Figure 2H) shows considerable conservation from algae to humans. Many key glycolytic and TCA cycle enzymes showing RNA-binding function are common across algae, plants, and humans. On the other hand, some moonlighting functions of a few enzymes are conserved between humans and algae, which is absent in plants. We found an exceptionally high (100%) coverage of creRBPs in the oxidative phosphorylation pathway. In plants, this is 57% (4 out of 7), and in humans, this is 85% (6 out of 7) (Figure 2I). Approximately 67% of the enzymes involved in fatty acid biosynthesis exhibited RNA-binding activity in our study, a proportion comparable to that observed in humans, but notably lower in plants. Our analysis revealed a high association of creRBPs with amino acid biosynthetic pathway enzymes (Supplementary Figure S2B and S2C). The coverage of creRBPs associated with the amino acid biosynthesis pathway is higher than that of RBPs identified from plants (Figure 2C). Extensive research has been conducted on the enzyme SHMT1, a key component of amino acid metabolism, particularly in the context of its RNA-binding properties and role in riboregulation. SHMT1 is a key enzyme required for the interconversion of glycine to serine. The riboregulation of SHMT1 by binding to RNA has been elucidated functionally and structurally^38^. We found 3 SHMTs in our creRBP and PBR datasets. It is quite evident that the RNA-binding function of SHMT1 is conserved in algae, which strongly suggests the preservation of its riboregulation. In this context, our finding of RNA binding activity of several amino acid biosynthetic enzymes suggests riboregulation might be a prevalent mode of fine tuning amino acid metabolism, which is an interesting area for future investigation.

### RNA binding activity of metabolic enzymes varies with the circadian time

To capture the temporal patterns of the RNA interactome across the circadian day-night cycle, cells were collected and processed from 3 independently grown cultures at LL48, LL54, LL60, and LL66, corresponding to subjective dawn, midday, dusk, and midnight, respectively (Figure 1A). The intensity of proteins of 518 creRBPs from all 12 replicates were normally distributed. (Supplementary Figure S3A). We compared the intra- and inter-time point correlation of the replicates (Supplemental Figure S3B). We found maximum variation in the subjective day vs subjective night time points. Further, we compared the variation in all replicates from subjective day and night (Supplementary Figure S3C), revealing the overall high reproducibility within replicates. Correlation heatmaps (Figure 3A) and principal component analysis (PCA) (Supplementary Figure S4A) demonstrated strong clustering by time point and high reproducibility among biological replicates. A clear distinction between day and night was evident in the RNA-binding activity of the creRBPs. In photosynthetic organisms, transitions between light and dark are critical and are often accompanied by vital changes in RNA and protein abundance. It is therefore reasonable to expect that rhythmic RNA-binding activity would exhibit a similar temporal pattern. Overall, minimal variation in RNA-binding activity was observed between the two daytime points (subjective dawn and subjective midday) or between the two nighttime points (subjective dusk and subjective midnight).

**Figure 3:**
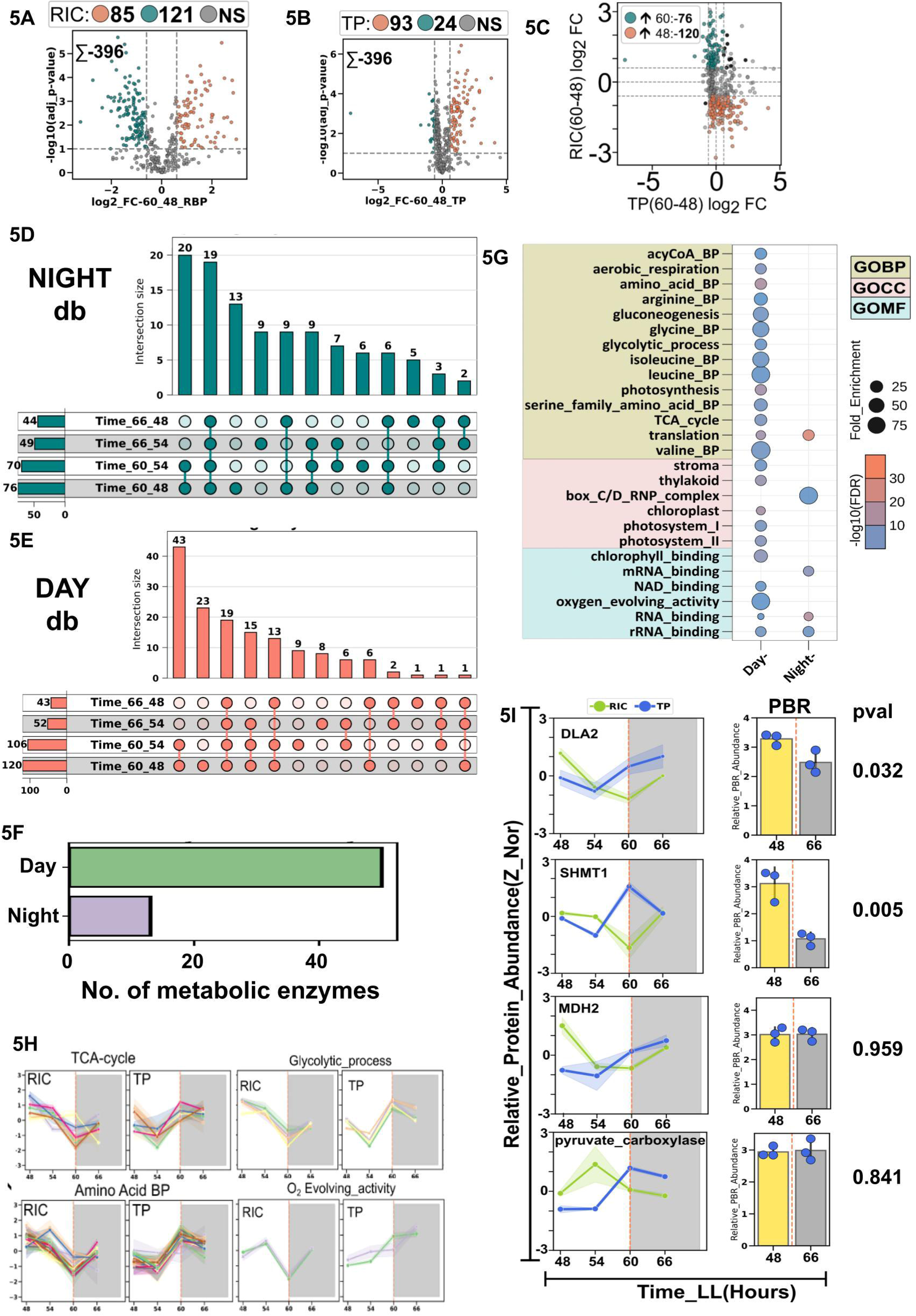
Temporal profiling of the RBP-RNA-Interactome in *C. reinhardtii*: **3A)** Heatmap depicting Pearson correlation coefficients among biological replicates across of subjective day and subjective night time points of the RIC experiment. Correlations were computed using Python. The color gradient represents the strength of correlation, with blue indicating high correlation and red indicating low correlation. **3B)** Pie chart represents the percentage of RPBs that exhibit temporal dynamicity, where teal and olive colors representing the percentage of cycling protein and non-cycling proteins respectively. **3C)** Heatmap depicting the distribution of day and night RBP (RNA-binding protein) clusters. **Top:** Night cluster, showing RBPs with elevated abundance during nighttime time points. **Bottom:** Day cluster, representing RBPs with higher abundance during daytime time points. In both panels, the X-axis denotes the time points, while the Y-axis represents the *z*-normalized relative protein intensity. The yellow line indicates the median intensity across each time point. color intensity reflects the number of RBPs: darker shades correspond to a higher number of RBPs. **3D)** Scatterplot representing the GO enrichment of the RBPs whose abundance in the interactome are higher during the day or the night. The fold enrichment and the -log10 FDR values are denoted by the diameter and color coding respectively. **3E)** Line plots illustrating the relative intensity of the proteins (enzymes) associated with metabolic pathways. The shaded areas represent the standard error mean. The grey-shaded rectangle represents the subjective night. The X- and Y-axis represents time and the relative protein intensity respectively. The yellow line represents the median. **3F-G)** Immunoblot showing the temporal recovery of ADH (3F: Left panel) and ATPA (3G) in the RIC samples across four circadian time points. Silver staining of the same blot is presented as a loading control. The accompanying histogram displays the relative quantification of ADH signal intensity from the immunoblot, while the point plot represents the relative intensity from the corresponding silver-stained lanes. The blue line represents the relative quantification of the silver staining. Line plot depicting the temporal profile of ADH and ATP A protein abundance obtained from SWATH-MS analysis across circadian time points. The gray shaded area indicates the subjective night. The X-axis denotes time, and the Y-axis represents relative protein intensity. **3H)** Scatter plot illustrating Gene Ontology (GO) enrichment analysis of Protein-Bound RNAs (PBRs) identified in the RIC experiment. Each dot represents a significantly enriched GO term. The size of the dots corresponds to the fold enrichment, while the color gradient reflects the statistical significance, represented as –log₁₀(FDR). **3I)** Hierarchical clustering-based heat maps with the z normalized FPKM values of the significantly varying PBRs across the subjective day-night samples.

Of the total 518 creRBPs identified, 372 (∼70%) exhibited significant temporal variation (Figure 3B). Notably, 109 (∼29% of total creRBPs) of these 372 creRBPs (Supplementary Figure S2D) were classified as metabolic enzymes, accounting for approximately 78% (109/140) of all identified metabolic enzymes with RNA-binding activity in *C. reinhardtii*. To further explore this, we generated a density plot of the 372 temporally regulated creRBPs. Two distinct clusters emerged: one comprising 200 proteins with enhanced RNA-binding activity during the day, and another of 172 proteins with higher RNA-binding at night (Figure 3C and Supplementary Figure S4B). The amplitude of the RBP-RNA interaction was higher (Supplementary Figure S4C) than what we got for the rhythmic proteins in our previous study^17^. Functional enrichment analysis revealed that day-active creRBPs were associated with the TCA cycle, ribosome assembly, photosynthetic electron transport, chlorophyll binding, aerobic respiration, amino acid biosynthesis, and oxidoreductive processes. Interestingly, chloroplast- and mitochondria-localized creRBPs exhibited both day- and night-specific activity, suggesting compartment-specific temporal regulation (Figure 3D).

More than 20% (83 out of 372) of the creRBPs exhibiting subjective day-night dynamicity in RNA-binding were derived from chloroplasts and mitochondria, exhibiting over 3-fold enrichment with a false discovery rate (FDR) below 1E-08. Functional enrichment analysis revealed greater than 25-fold enrichment in components of the photosynthetic electron transport chain and over 35-fold enrichment in Calvin cycle enzymes. Similarly, mitochondrial processes, including oxidative phosphorylation, aerobic respiration, and the tricarboxylic acid (TCA) cycle, each displayed approximately 10-fold or higher enrichment, with statistically significant FDR values. To temporally map RNA-binding activity of proteins and enzymes involved in key metabolic pathways, we plotted their RNA-binding capacities over time. Notably, RNA-binding activities associated with mRNA processing, box C/D small nucleolar RNP (snoRNP) complexes, and mitochondrial ribosomes were predominantly observed during the subjective night. In the TCA cycle, fatty acid biosynthesis, amino acid biosynthesis, the electron transport chain, and aerobic respiration, the RNA-binding pattern of associated proteins were highly coordinated, showing a marked decline during subjective dusk. An exception was observed in the box C/D snoRNP complex, which showed increased RNA-binding activity at subjective dusk (Figure 3E). We found some creRBPs associated with mitochondria, chloroplasts, translation, and photosynthesis are preferentially binding RNA during the subjective day, while other creRBPs from the same prefer subjective night for RNA-binding (Supplementary Figure S4D). To validate the temporal dynamics of RNA-binding revealed by SWATH-MS, we selected two representative metabolic enzymes, ATP synthase subunit alpha (ATPA) and alcohol dehydrogenase (ADH), both of which exhibited time-dependent RNA-binding. Using immunoblotting of circadian RIC samples, we confirmed the temporal RNA-binding patterns of ATPA and ADH, corroborating with our findings from the SWATH-MS analysis (Figure 3F and 3G). These findings strongly suggest that circadian RNA-binding to metabolic enzymes may modulate their activity, thereby influencing corresponding metabolic processes. Our study identified several key enzymes from different cellular metabolic pathways showing robust temporal regulation in RNA-binding, suggesting a widespread involvement of circadian riboregulation in orchestrating daily rhythms of metabolic activity.

Do the captured PBRs from the RBP–RNA interactome also exhibit day–night dynamics? To address this, we focused on the temporal binding affinities of mRNAs to their cognate creRBPs (Figure 1A). The observed temporal enrichment of specific mRNAs may reflect either a time-dependent preference for their translation or their involvement in circadian riboregulation. Previous research reported the possibility of both mechanisms in tandem that fine tunes cellular metabolic outputs. Biological triplicates from subjective day and night samples clustered distinctly, indicating a clear temporal pattern in mRNA binding by creRBPs (Supplementary Figure S5A). Pairwise correlation analysis further confirmed strong concordance among replicates (Supplementary Figure S5B). From this dataset, we identified 87 and 101 mRNAs with preferential binding to their cognate creRBPs during the subjective day and night, respectively (Figure 3H and Supplementary Figure S5C). Functional enrichment analysis of the day-binding mRNAs revealed significant overrepresentation of processes related to photosynthesis, chloroplast function, and Photosystem I and II, consistent with our previous findings from circadian proteomics^17^ (Figure 3I). This suggests that the enrichment of photosynthesis-associated mRNAs during the day may support time-aligned protein synthesis that prefer subjective day. In contrast, 101 mRNAs were enriched during the night and we found an enrichment of metabolic processes, carbohydrate metabolism and peroxisomes. We found 385 creRBPs whose mRNA is also present in our captured PBRs (Supplementary Figure S5D). They might represent feedback regulation, riboregulation by binding to other creRBPs or conventional regulation of synthesis of their proteins, which requires further investigation.

### RNA binding activity metabolic enzymes is independent of their abundance profile across circadian time

The next obvious question that we set out to address was whether the temporal RNA-binding pattern of the creRBPs matches with their abundance profile. For this we took advantage of our earlier circadian proteome study with *C. reinhardtii* cells^17^. The cells used for total proteome and the current RIC study were grown under the same conditions and captured at the same time points (LL48, LL54, LL60 and LL66). Comparing the creRBPs apprehended from RIC with the total proteome, we found 396 common creRBPs (Figure 4A). Using these 396 creRBPs we compared whether their abundance concurred with their RNA-binding profiles. To assess the overall correlation between protein abundance and RNA-binding, we performed regression analysis by plotting the log2 intensities of creRBP abundance in the RIC (Y-axis) against their corresponding values in the total proteome (X-axis) at each circadian time point. A higher R² value would indicate a positive correlation between RNA-binding activity and protein abundance, suggesting that RNA-binding is primarily driven by protein abundance. The highest R² value observed was 0.32 at subjective midnight, and this weak correlation persisted across all circadian time points (Figure 4B), indicating a general disagreement between creRBP abundance and their RNA-binding activity. RNA-binding dynamicity independent of their protein levels has been previously reported in drosophila embryos when early and late developmental stages were compared^77^ and during cardiac growth^78^.

**Figure 4:**
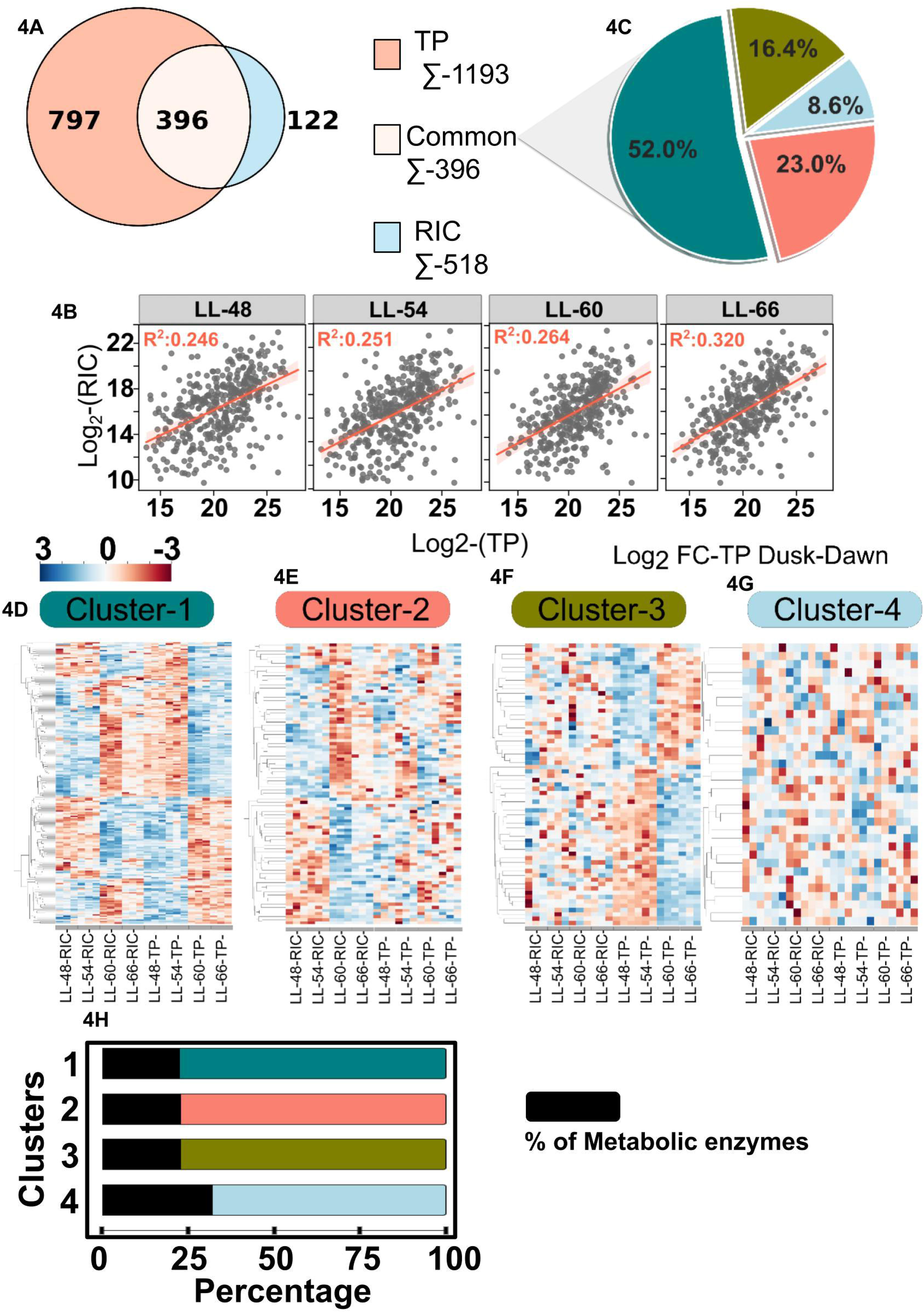
creRBPs exhibit RNA-binding independent of their abundance in the proteome: **4A)** Venn diagram illustrating the overlap and unique proteins identified in the RIC dataset compared to the total proteome. **4B)** Scatterplot showing the correlation between average log₂ protein intensities in the RIC and total proteome across four time points. The orange line indicates the regression fit, with corresponding R² values. The X-axis denotes the log₂-transformed relative protein intensities in the total proteome (TP, *n* = 3), while the Y-axis represents the log₂-transformed relative protein intensities in the RIC dataset (*n* = 3). The respectctive timepoints and their R2 values are indicated on the top of each panel. **4C)** Pie chart illustrating the distribution of the common 396 RBPs across four distinct clusters, which are defined based on their temporal cycling patterns observed in the RIC and total proteome datasets. Each cluster represents a group of RBPs with their respective abundance profiles in RIC and TP described in the heatmaps below. **4D-G)** Hierarchically clustered heatmaps illustrating the abundance profiles of RBPs in both the RIC and total proteome datasets across four circadian timepoints, categorized according to the clusters defined in the preceding pie chart. Each heatmap corresponds to one of the four clusters and displays the temporal abundance dynamics of RBPs captured in the RIC or total proteome. The color scale represents Z-score–normalized protein intensities, allowing for comparative visualization of relative abundance patterns. The X-axis indicates the circadian timepoints for both RIC and TP conditions, enabling a direct comparison of RNA-binding dynamics versus total protein abundance. **4H)** A bar plot illustrating the distribution of metabolic enzymes across the four defined clusters in the preceding pie chart and the hierarchically clustered heatmaps.

We further categorized the 396 creRBPs into four distinct clusters based on their abundance profiles in total proteome and RIC. The first cluster, comprising 52% of the creRBPs, includes proteins whose abundance and RNA-binding activity both exhibit clear temporal variation (Figure 4D). However, their dynamic patterns appear to differ across circadian time points (Figure 4D). The fourth cluster, the smallest, includes approximately 9% of the creRBPs, which show no significant changes in either abundance or RNA-binding activity across the 24-hour circadian cycle (Figure 4G). The second cluster contains 23% of the creRBPs, characterized by temporally dynamic RNA-binding activity despite consistent protein abundance across the circadian cycle (Figure 4E). In contrast, the third cluster, encompassing 16% of the creRBPs, exhibits temporal changes in protein abundance without corresponding changes in RNA-binding patterns across circadian time points (Figure 4F). These findings reveal a widespread incongruence between the temporal regulation of RNA-binding activity and protein abundance of the RBPs, indicating that RNA-binding dynamics are frequently uncoupled from changes at the protein levels. Of the 396 creRBPs, approximately one-quarter (94 proteins) are metabolic enzymes. Each of the four clusters contains at least ∼20% representation from metabolic enzymes. (Figure 4H) We propose that a substantial number of these metabolic enzymes may function as “circadian dynamic binders,” wherein temporal patterns of RNA-binding activity are independent of their respective protein abundance. Temporal classification and functional enrichment analysis of these metabolic enzymes may reveal biologically significant trends and regulatory mechanisms underlying circadian metabolic control.

### Metabolic enzymes as circadian dynamic binders suggesting a widespread role of riboregulation in temporal alteration of metabolism

Do circadian dynamic RNA binders exhibit a distinct pattern of functional enrichment between the subjective day and night? To address this, we used a pairwise comparison approach between all possible subjective day vs night combinations (i.e., subjective midnight vs subjective midday, subjective midnight vs subjective dawn, subjective dusk vs subjective midday, subjective dusk vs subjective dawn) to ascertain the time-of-the-day dependent abundance and RNA-binding profiles in the total proteome and RIC respectively. For each of these pairwise combinations, we first selected only those creRBPs that showed clear and significant subjective night vs day differences using a standard cutoff of 1.5 fold change and -log10 (adjusted p value) of 1(Figure 5A and 4B) in their abundance and RNA-binding profiles (Supplementary Figure S6A). We then merged the separate abundance vs RNA-binding volcano plots into a single X-Y coordinate plots to extract the subjective day-dynamic and subjective night-dynamic creRBPs (Figure 5C and Supplementary Figure S6B). We realized that due to the pairwise comparisons, circadian dynamic creRBPs may appear redundantly across multiple time point combinations, such as LL60 Vs LL48, LL66 Vs LL48, LL60 Vs LL54, and LL66 Vs LL54. To address this, we separated and mapped the redundant and unique dynamic binders’ specific to the subjective day and night phases (Figure 5D, E). In total, we identified 147 and 108 creRBPs that preferentially bind RNAs during the subjective day and night, respectively, which misaligns with their protein levels (Figure 5D E). Overall, about 50% (255/518) of the total creRBPs qualified as circadian dynamic binders, and 63 of those are metabolic enzymes (Figure 5F). Among the 255 dynamic RNA-binding proteins (RBPs), approximately one-quarter are associated with chloroplast and mitochondrial proteins and enzymes. These also include enzymes involved in the metabolism of amino acids such as glycine, valine, isoleucine, serine, and leucine. Notably, several enzymes central to cellular energetics, including those from the TCA cycle, glycolysis/gluconeogenesis, and the pentose phosphate pathway, exhibit preferential RNA binding during the subjective day, despite their protein abundances typically peaking at subjective night (Figure 5G). To highlight the difference between the dynamicity of RNA-binding and total protein abundance, we specifically selected enzymes and proteins from key metabolic pathways and analyzed their respective profiles (Figure 5H and Supplementary figure S7A). Additionally, several enzymes involved in amino acid biosynthesis also exhibit subjective day preference for RNA binding, further emphasizing the circadian regulation of core metabolic functions through riboregulation. The temporal RNA-binding patterns observed in many metabolic enzymes, which occur independently of changes in their protein abundance, warrant further investigation to uncover the underlying mechanisms driving this circadian binding behavior.

**Figure 5:**
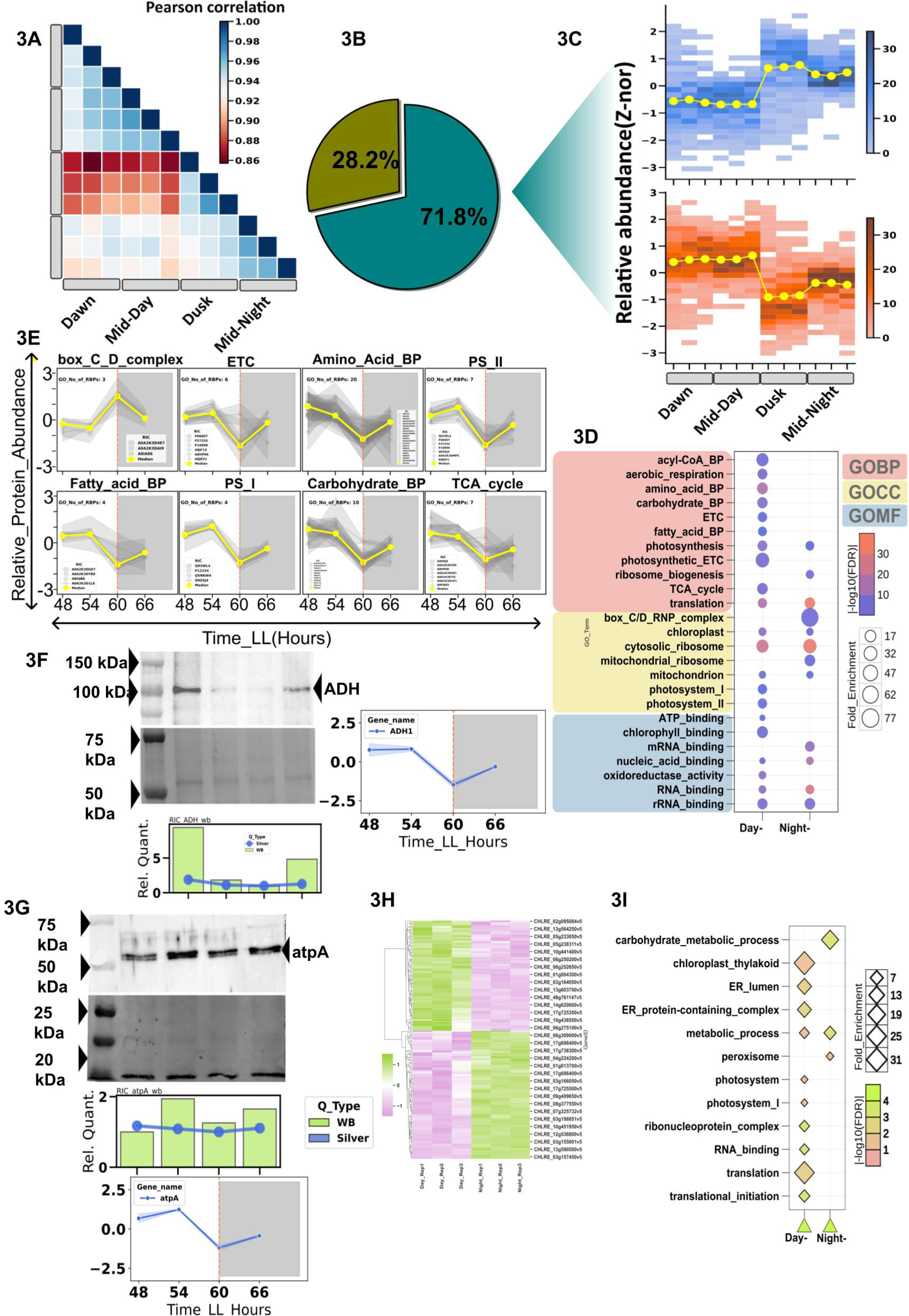
Circadian RBPs are dynamically binding with the RNA: **5A-B)** Volcano plots depicting the differential protein abundance between subjective dusk (LL60) and subjective dawn (LL48) in the RIC (A) and the total proteome (TP) (B). The X-axis represents the mean log₂ fold change, while the Y-axis represents the –log₁₀ of the adjusted p-value. Proteins significantly upregulated at subjective dusk (log₂ fold change ≥ 0.6 and –log₁₀ adjusted p-value > 1) are highlighted in orange, whereas proteins significantly upregulated at subjective dawn (log₂ fold change ≤ –0.6 and –log₁₀ adjusted p-value > 1) are shown in teal. The vertical and horizontal dashed lines indicate the fold change and significance thresholds, respectively. **5C)** To portray the dynamicity (where RNA binding function of the creRBPs do not correspond to their abundance profiles from TP) we show a volcano plot. The Y-axis represents the log₂ fold change in mean protein abundance between LL60 Vs LL48 in the RIC, while the X-axis represents the corresponding change in the total proteome (TP). Light grey dots indicate proteins with no significant change in either dataset. Black dots represent proteins with concordant and significant changes in both RIC and TP, suggesting abundance-driven RNA binding. Teal-colored dots denote “dynamic binders” upregulated at subjective dusk, which represents proteins with significant enrichment in the RIC but not in the TP. Conversely, orange dots represent “dynamic binders” enriched at subjective dawn, also showing RIC-specific regulation independent of total abundance. The vertical and horizontal dashed lines indicate the fold change and significance thresholds, respectively. **5D-E)** A upset plot representing the common and unique “dynamic Binders” at subjective night and day respectively. **5F)** Histogram representing the number of metabolic enzymes in the day and night dynamic binders. **5G)** Gene ontology (biological process, cellular component and molecular function) enrichment of the day and night dynamic binders. **5H)** Line plots illustrating the abundance profiles of enzymes identified as creRBPs from selected metabolic pathways, measured across circadian time points in both the RIC and TP datasets, highlighting key enzymes that are “circadian dynamic binders”. **5I) 1st panel-left:** Line plot representing the abundance profile of the metabolic enzyme dihydrolipoamide acetyltransferase (DLA2) of the pyruvate dehydrogenase complex in the RIC (green line) and TP (blue line) across the circadian day and night time points. The shaded area represents the +/-1 SEM. **1st panel-right:** Histogram representing the relative abundance of DLA2 mRNA in the PBR fraction at day and night. The DLA2 mRNA is significantly changing in day and night. The yellow and grey bar represents subjective day and night respectively. The three replicates from day and night are color as blue circles. **2nd panel-left:** Line plot showing the dynamic binding of the metabolic enzyme SHMT1. The color scheme is same as above. **2nd panel-right:** Histogram showing the relative abundance of SHMT1 mRNA in the PBR fraction at day and night. The SHMT1 mRNA is significantly changing at day and night. **3rd panel-left:** Line plot illustrating the dynamic binding of the metabolic enzyme MDH2. color scheme is same as above. **3rd panel-right:** Histogram showing the relative abundance of MDH2 mRNA in the PBR fraction at subjective day and night. The mRNA of MDH2 is not significantly different at day and night. **4th panel-left:** Line plot illustrating the dynamic binding of the metabolic enzyme PC. color scheme is same as above. **4th panel-right:** Histogram showing the relative abundance of PC mRNA in the PBR fraction at subjective day and night. The mRNA of PC is not significantly different at day and night.

Interestingly, from previous literature we identified riboregulatory functions for five different metabolic enzymes, four from mammals^34,35,38,39^ and one from *C. reinhardtii*^67^. SHMT1, an enzyme involved in amino acid metabolism, has been shown to moonlight as an RNA-binding protein (RBP) by binding to and regulating the translation of SHMT2^38^. This interaction, in turn, modulates SHMT1’s own enzymatic activity. In *C. reinhardtii*, we identified at least three SHMT isoforms, two of which exhibit RNA-binding moonlighting functions. Both show time-of-the -day dependent variation in RNA-binding capacity, with enhanced binding during the subjective day (Figure 5I, 2^nd^ panel left and Supplementary Figure S7B top left). On the other hand we found all three SHMTs in the PBR pool, an observation that is similar to mammalian SHMTs thereby demonstrating a mammalian-like functional conservation. Notably, SHMTs were more abundant in the PBR during the day, supporting the idea of multiple regulatory layers for fine-tuning metabolic outputs (Figure 5I, 2^nd^ panel right and Supplementary Figure S7B top right). Similarly, the TCA cycle enzyme MDH2 has been shown in mammals to bind various coding and non-coding RNAs, competing with NAD⁺ for binding and thus modulating its enzymatic function^35^. In *C. reinhardtii*, we identified two MDH isoforms, cytosolic and chloroplastic, both of which exhibit RNA-binding moonlighting functions. These isoforms also display a preference for RNA-binding during the subjective day (Figure 5I, 3^rd^ panel left and Supplementary Figure S7B bottom left). While MDHs were detected in the PBR, no dynamic changes in their levels were observed over the subjective day-night cycle (Figure 5I, 3^rd^ panel right and Supplementary Figure S7B bottom right). In the case of DLA2, a component of the pyruvate dehydrogenase complex in *C. reinhardtii*, previous studies have reported RNA binding upon acetylation^67^. Our findings confirm that DLA2 binds RNA and further reveal a clear subjective day-night preference in its RNA-binding activity (Figure 5I, 1^st^ panel left). DLA2 was also detected in the PBR, with significantly higher abundance during the subjective day compared to the subjective night (Figure 5I, 1^st^ panel right). Pyruvate carboxylase (PC) is a key enzyme that replenishes the TCA, initiates gluconeogenesis and plays a key role in lipid metabolism^79^. RNA-binding has been shown to regulate the activity of pyruvate carboxylase (PC) and influence lipid metabolism in mammals^39^. In our study, PC emerged as a dynamic creRBP, exhibiting a subjective day preference in its RNA-binding activity, independent of its protein abundance (Figure 5I, 4^th^ panel left). While PC mRNA is present in the PBR pool, it does not show temporal dynamicity (Figure 5I, 4^th^ panel right). This makes PC a compelling candidate for exploring its role in circadian lipid metabolism, a feature of particular interest for biofuel research. These enzymes represent potential targets of circadian riboregulation that may ultimately contribute to the temporal regulation of the metabolic processes they are involved in.

Our cellular energetics map integrates glycolysis, the TCA cycle, the pentose phosphate pathway (PPP), and related metabolic networks. Our study reveals that many components of these three pathways function as subjective day dynamic RNA binders (Figure 6A), suggesting a coordinated regulatory mechanism in which RNA-binding can modulate their activity. Glycolysis, TCA cycle and PPP are interconnected pathways that form the central backbone of cellular metabolic homeostasis. Circadian riboregulation of enzymes involved in these pathways presents a compelling avenue for further investigation to better understand the daily regulation of metabolism. In autotrophs, photosynthesis serves as a major energy-generating pathway and is also a primary source of atmospheric oxygen. Light harvesting is a critical component of this process, fuelling carbon fixation. Our data show that numerous enzymes and proteins associated with the photosynthetic apparatus, including light-harvesting complexes, photosystem I and II proteins, and components of the oxygen-evolving complex, display RNA-binding activity preferentially during the subjective day (Figure 6B), which does not depend on their protein abundance. To validate the SWATH proteomics results in the context of circadian dynamic binders, we used an alternative approach of immunoblotting. We chose a circadian dynamic binder, in this case *psbD,* whose antibody is commercially available. In our proteomic analysis, we found the RNA-binding activity of *psbD* is high during the subjective day, lowest during the subjective dusk, and increases again by the subjective midnight. The total protein abundance of *psbD* is low during subjective day but high during subjective night. Although the protein abundance increases during day to night transition, the RNA-binding activity decreases at the transition before regaining the activity later during the night. Our immunoblotting study with captured interactome and total proteome at equivalent time points concurs with our conclusions drawn from the SWATH-MS (Figure 6C). It is known from earlier studies^80^ that *psbD* protein shows reversible phosphorylation under LL conditions. We envisage that *psbD* clock driven phosphorylation/ dephosphorylation dynamics might play a role to regulate its RNA-binding. Earlier studies have shown that phosphorylation/dephosphorylation can regulate the RNA binding activity of proteins^81–83^.

**Figure 6:**
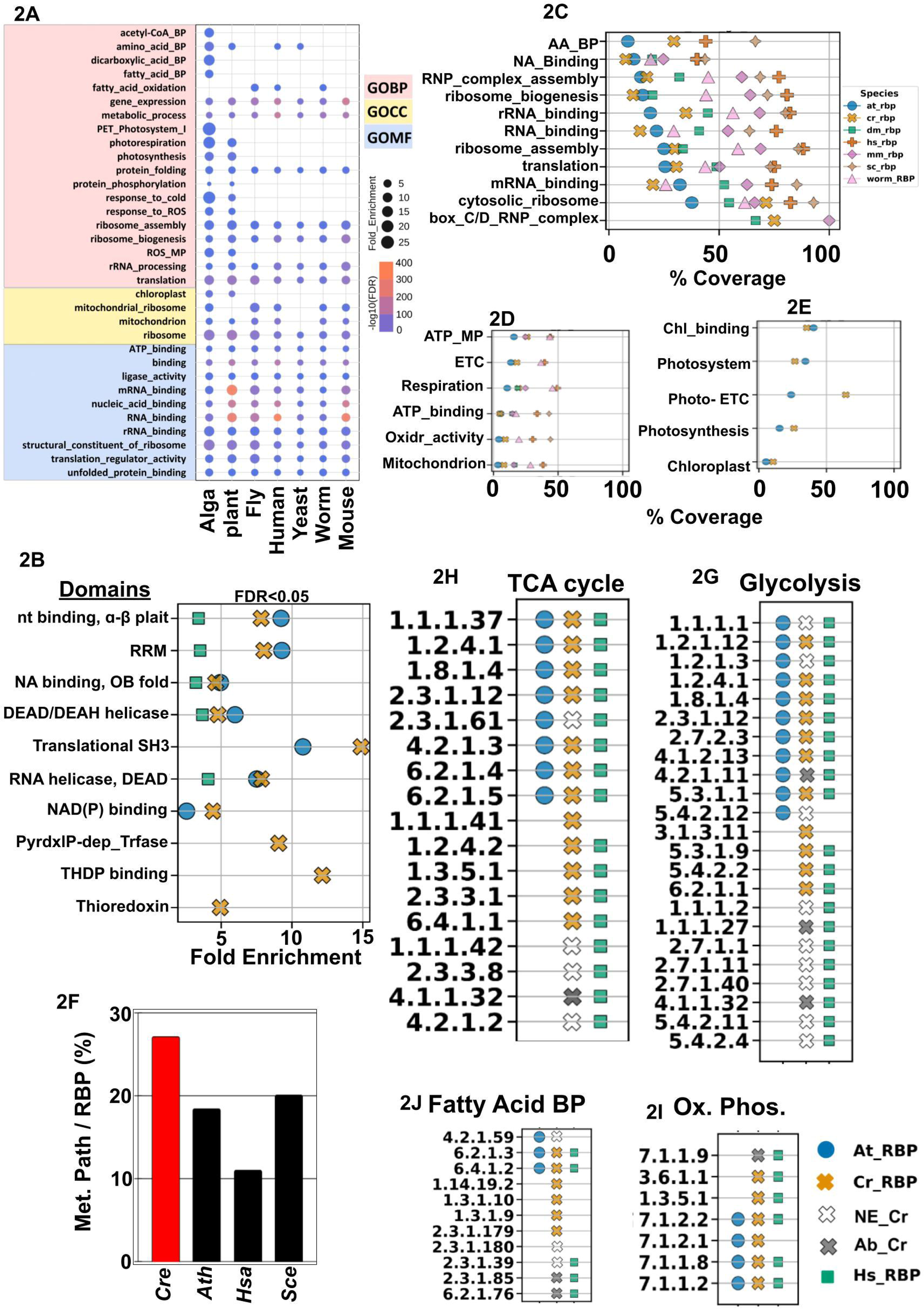
Many dynamic binders are associated with key physiological and metabolic processes: **6A)** The schematic illustrates a segment of the cellular energetics pathway, including glycolysis, the TCA cycle, and the pentose phosphate pathway. The enzymes associated with the specific reactions are denoted by their EC number. Many enzymes in these pathways are enriched as dynamic binders among the identified creRBPs. Dynamic creRBPs active during the subjective day are shown in blue, those active at night in yellow, and those with dynamic binding at both times are depicted as half blue and half yellow. **6B)** Schematic representation of the RNA-binding proteins in the chloroplast. The diagram illustrates the day dynamic creRBPs associated with photosynthesis, which is another major energy generating organelle in photoautotrophs. Here specifically shown the creRBPs involved in the Light Harvesting Complex, Photosystem I and II localized within the thylakoid membrane. **6C)** Validation of day-specific dynamic RNA binding to psbD, identified as a creRBP target. The left panel illustrates the circadian expression profile of psbD at the protein level. From top to bottom: an immunoblot showing psbD protein levels across four circadian time points; a silver-stained version of the same blot used as a loading control; a histogram displaying the relative quantification of psbD band intensities from the immunoblot, overlaid with line plots representing the corresponding silver staining intensities; and a line plot depicting the relative abundance of psbD protein measured in the total proteome using SWATH-MS. The right panel shows *psbD* abundance in the circadian RNA interactome. The sequence of figures, experiments, and analyses mirrors that of the left panel, with one key difference: instead of the total proteome, RNA-bound proteins captured by RIC across four circadian time points were analyzed. Additionally, the silver-stained gel in this panel uses the RNase band as the loading control. **6D) Left:** Schematic representation of the dynamic RNA-binding proteins (RBPs) associated with the box C/D pathway. Colored proteins indicate RNA-binding activity, while red and green arrows represent a decrease and increase in RNA-binding, respectively. The observed nighttime increase in RNA-binding may influence ribosome biogenesis. **Right:** Schematic representation of the endoplasmic reticulum (ER) highlighting RNA-binding proteins (RBPs) localized within it. These RBPs are involved in protein processing in the ER and exhibit dynamic binding patterns at night, potentially influencing the efficiency of protein processing.

NOP56 and NOP58 proteins are members of the Box C/D RNP complex within the nucleolus region. NOP56/NOP58 are a part of the snoRNA-protein complex that regulates ribosome biogenesis and the 2’-O-methylation of coding and non-coding RNAs^84^. Our study shows preferential binding of NOP56 and NOP58 at night, suggesting a possible mechanism that can regulate the RNA-methylation machinery temporally (Figure 6D). We found the ER-localized protein disulfide isomerase (PDI) to bind RNA with a temporal preference. Its RNA-binding activity is more pronounced during the night (Figure 6D). A chloroplast-localized PDI has been previously shown to regulate the synthesis *psbD* protein by binding to the 5’ UTR of its mRNAs in a redox responsive manner^85,86^. One sub-family of PDI has been shown to regulate circadian rhythms of phototaxis in *C. reinhardtii*^87^.

## Discussion

It is a well-known fact that many metabolic enzymes possess RNA-binding as a ‘moonlighting’ function^31,32^, where interaction with cognate RNAs can modulate their enzymatic activity and the efficiency of their associated metabolic pathways, ultimately leading to distinct physiological outcomes^34,39^. RBP–RNA interactions can play complex roles, as exemplified by studies on SHMT1^38^. SHMT1 functions as a conventional RBP by binding to the 5′UTRs of its own mRNA and that of SHMT2, thereby modulating SHMT2 translation. Simultaneously, this RNA interaction also regulates SHMT1’s enzymatic activity, thereby regulating the cellular glycine-serine balance^38^. In our datasets we also found 3 isoforms of SHMTs, both in the creRBP and PBR fractions, which align to its functional conservation across species. However, whether global cellular RBP–RNA interactions have a wider impact on overt circadian metabolic and physiological processes remains an open question. To address this overarching question, we first sought to determine whether RNA–RBP interactions exhibit temporal dynamics over the daily cycle. We believe this fundamental approach will enhance our understanding of circadian riboregulation and its potential role in orchestrating daily metabolic rhythms.

To ensure that the observed circadian dynamics reflected true endogenous rhythms and not residual effects of prior light-dark (LD) cycles, cells entrained under LD conditions were transferred to constant light (LL) for 48 hours prior to sample collection. Although free-running conditions can cause damping of the rhythmic amplitude^88–91^, this approach was accepted as a necessary trade-off to capture circadian dynamicity. Our study reveals that 72% (372 out of 518) of *C. reinhardtii* RBPs (creRBPs) exhibit temporal changes in RNA-binding between the subjective day and night, with approximately one-third associated with metabolic processes. To our knowledge, this is the first comprehensive profiling of temporal RNA interactome dynamics across multiple circadian time points, highlighting the potential for circadian riboregulation to drive daily metabolic rhythms. Among these, 255 creRBPs were identified as circadian dynamic binders, showing rhythmic RNA-binding patterns that are uncoupled from their total protein abundance, suggesting regulation directly by the internal circadian clock rather than through their protein levels, which is also clock controlled^17^. This phenomenon of temporally dynamic RNA–RBP interactions has been observed in diverse biological contexts, such as development^77^, cardiac growth^78^, and viral infections^92^. Our data extend this concept to circadian regulation, identifying RBPs that bind RNA with temporal dynamicity and can act as potential drivers of rhythmic metabolic outputs through riboregulation. Enrichment analysis indicates widespread localization of creRBPs to chloroplasts and mitochondria, where many display circadian RNA-binding dynamics. We propose that moonlighting RNA-binding functions of metabolic enzymes in these organelles play a critical role in coordinating daily metabolic rhythms. Given that the Calvin cycle is the primary biochemical pathway for fixing atmospheric CO₂ into carbohydrates, the RNA-binding dynamicity of its enzymes presents a promising avenue for exploring the circadian regulation of carbon fixation via circadian riboregulation. Mitochondria, as the central site of oxidative phosphorylation, also play critical roles in lipid metabolism and reactive oxygen species (ROS) regulation^93^. Their involvement in circadian metabolic regulation is well established^94^. Our study not only reinforces this well-studied paradigm but also identifies novel candidate targets, suggesting that riboregulation of key mitochondrial enzymes may be a valuable area for further investigation of metabolic homeostasis across the daily cycle.

One plausible explanation for the extensive circadian dynamics in the RNA-binding moonlighting functions of many metabolic enzymes might be the involvement of specific post-translational modifications^34,67^. These modifications can either promote RNA-binding, thereby altering the enzymes’ primary catalytic activities or can induce a functional shift from being an enzyme to a conventional RBP, warranting further investigation into the regulatory mechanisms underlying this functional duality. This has been demonstrated in the dihydrolipoamide acetyltransferase (DLA2) enzyme in *C. reinhardtii*, where acetylation of a single lysine regulates its functional transition from being a metabolic enzyme to an RBP^67^, and in enolase 1^34^, where RNA-binding regulates its catalytic function in glycolysis. In our datasets DLA2 exhibits temporal dynamicity with high RNA-binding observed during subjective day as compared to subjective night (Figure 5I). In mammals, acetylation of enolase1, a glycolytic enzyme, has been shown to increase its RNA-binding activity^34^. In this context, circadian acetylome studies in mouse liver have demonstrated that the acetylation of mitochondrial proteins exhibits a robust time-of-day–dependent regulation^25^. The most affected pathways include glycolysis/ gluconeogenesis, the TCA cycle, and fatty acid and amino acid metabolism^25^. Notably, the pathways enriched in the circadian acetylome show strong overlap with those identified in our enriched circadian RNA interactome, suggesting a close functional connection between these two regulatory layers.

Previous studies reported at least 5 RNA-binding metabolic enzymes involved in glycolysis, the TCA cycle, and amino acid metabolism not only exhibit altered enzymatic activity upon RNA binding, but also significantly impact cellular metabolic homeostasis and physiology. Interestingly, our study found that four of these five enzymes display temporal dynamics in their RNA-binding activity. Given their already crucial role in cellular metabolism, these enzymes represent key targets for investigating circadian riboregulation and its role in orchestrating temporal metabolic patterns. In addition, we discovered 109 metabolic enzymes that demonstrate time-of-the-day dependent dynamicity in RNA-binding and of these, 63 dynamically bind RNA across circadian cycles independent of their protein levels (Figure 3D, 4H and 5F). Finally, we propose a model illustrating how our study can serve as a foundational platform for understanding circadian regulation of metabolism through time-of-day-dependent riboregulation (Figure 7). We suggest that circadian regulation of post-translational modifications (PTMs), such as acetylation, may underlie the temporal RNA-binding activity of enzymes and proteins, thereby modulating their function to achieve circadian metabolic outputs. We believe our study will play a pivotal role in advancing our understanding of circadian riboregulation, a process wherein the circadian regulation of metabolic enzyme activity is mediated by their temporal dynamicity of RNA-binding function. These temporal regulatory mechanisms may represent a crucial piece in decoding the circadian patterns of cellular metabolism. Our findings lay the groundwork for a new understanding of temporal metabolism through the lens of circadian riboregulation.

**Figure 7:**
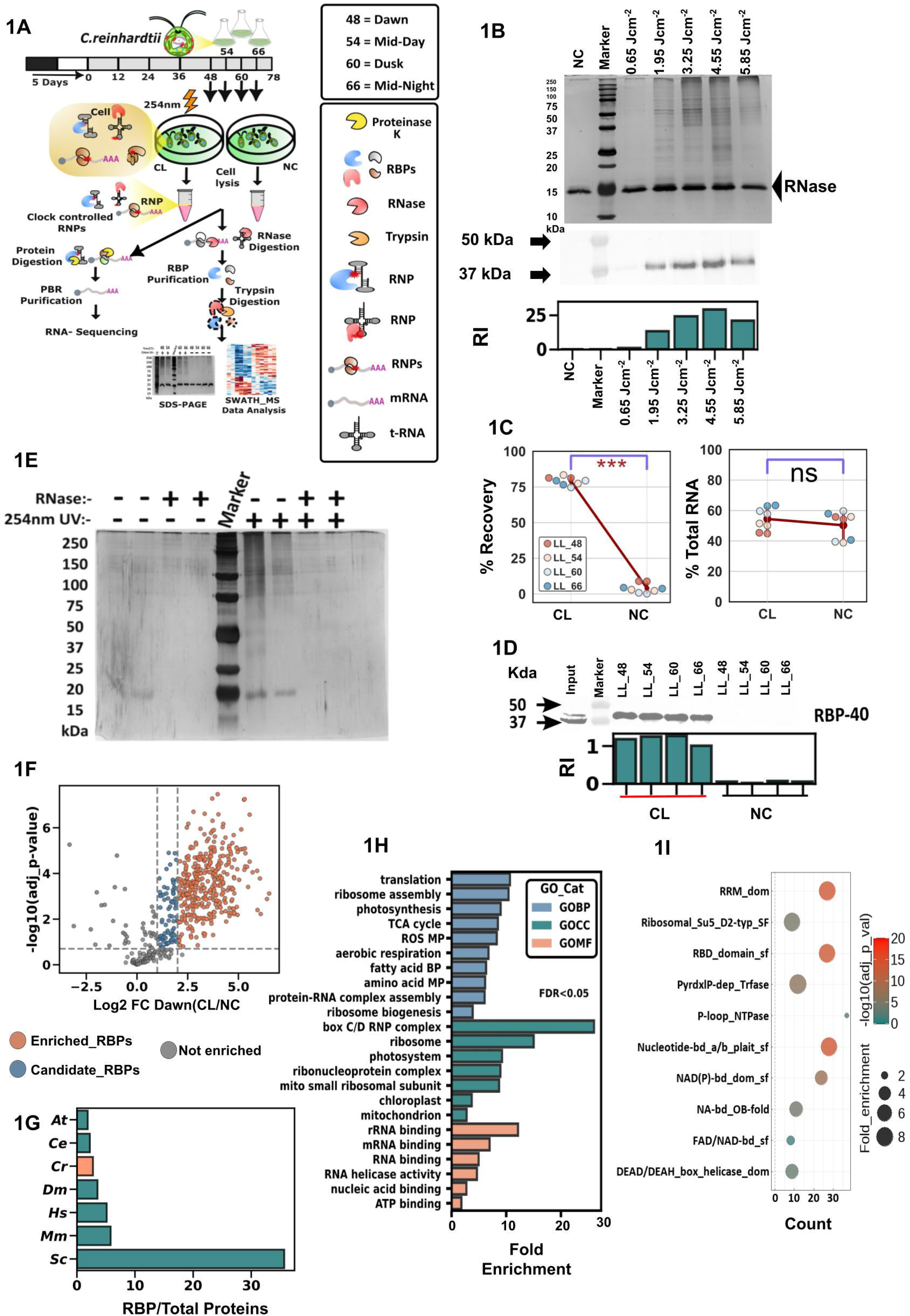
Modelling circadian riboregulation: Insights from temporal dynamics of RNA-binding and total proteome profiles. We propose a model illustrating how our findings lay the foundation for understanding circadian riboregulation and its role in the temporal control of cellular metabolism. We envision that metabolic enzymes exhibit time-of-day–specific shifts in their RNA-binding activity, contributing to the regulation of metabolic processes across the circadian cycle. While a complete global picture is yet to emerge, we believe that post-translational modifications (PTMs), such as acetylation, may play a critical role in modulating these temporal changes in RNA binding, potentially altering enzyme states (from enzyme state A to B) and influencing their RNA-binding affinity in a circadian manner. This circadian RNA-binding activity may, in turn, regulate the metabolic processes associated with the respective enzymes in a time-of-the-day dependent manner.

The types of RNA bound by different metabolic enzymes are notably distinct. For example, the glycolytic enzyme ENO1^34^ predominantly binds mRNAs, whereas the TCA cycle enzyme MDH2^35^ shows a preference for tRNAs. MDH2 contains an NAD (P) binding fold, and both NAD⁺ and tRNAs compete for this site. Since NAD⁺ is essential for MDH2 enzymatic activity, tRNA binding could potentially inhibit its function. In contrast, ENO1 lacks an NAD⁺ binding pocket, and previous studies have shown that it binds either its substrate or cognate mRNAs, but not both simultaneously. On the other hand, SHMT1 can bind to its own as well as to SHMT2 mRNA, which inhibits SHMT2 translational efficiency^38^, PC^39^ and p62^44^ binds to long and small noncoding RNAs respectively. To capture this diversity and comprehensively identify creRBPs regardless of their RNA-binding preferences, we employed an unbiased orthogonal phase separation approach. The identification of SHMTs in both the creRBP and PBR pools suggests a conserved functional role, similar to that observed in mammals. Overall, we found that the mRNAs of 385 out of 518 creRBPs are also present among the 2,476 captured PBRs (Supplementary Figure 5D). Whether this indicates a widespread feedback mechanism or suggests that their regulation is fine-tuned through multiple layers of post-transcriptional control remains to be determined through future studies. MDHs, PC and DLA2 not only exhibit RNA-binding activity but their corresponding mRNAs are also enriched in the PBRs (Figure 5I and S7B).

Notably, *C. reinhardtii* demonstrates a high degree of conservation in key metabolic enzymes that is often comparable to, or exceeds, that observed with humans rather than plants, highlighting its unique evolutionary position and ancestral links to both lineages (Figure 2 and S2). Furthermore, we demonstrated that many of these *C. reinhardtii* metabolic enzymes are circadian dynamic binders (Figure 4), and have also been recognized as RBPs across eukaryotes and in humans. Given their functional conservation across species, we propose that their riboregulatory role is likely preserved as well. Thus, the underlying mechanism governing riboregulation in these conserved enzymes may be identical in humans and *C. reinhardtii*, making *C. reinhardtii* a simple yet powerful model for further research in this context. Therefore, we propose that *C. reinhardtii* may serve as an effective and simple testbed for uncovering how the circadian clock regulates and coordinates multiple cellular metabolic processes through riboregulation, owing to its retention of both animal- and plant-like metabolic enzymes with moonlighting functions. Our findings underscore the pivotal role that temporal RNA-binding can play in modulating metabolic activities over the daily cycle. The extreme conservation of enzymes involved in crucial biological processes with moonlighting RNA-binding function suggests the deep-rooted existence of circadian riboregulation.

## Materials and Methods

### Cell Culture

Wild-type *Chlamydomonas reinhardtii (CC125)* were received as a kind gift from Prof. B. J. Rao of IISER Tirupati (originally obtained from Chlamycollection.org). CC125 cells were maintained in a 12:12 hours of light: dark regime at 18 ± 1 °C on TAP agar plates supplemented with 100ug/ml carbenicillin and 40 µg/ml Carbendazim in a Percival algal growth chamber (AL41L4) with 74 µmoles m^-2^ s^-1^. A loopful of *C. reinhardtii* cells were inoculated from 3 different plates in three different 100 ml Erlenmeyer flasks containing 40 ml of Tris-acetate-phosphate (TAP) liquid media plus 100 µg/ml carbenicillin and entrained under 12 hours of light and 12 hours of dark cycles for 5 days. Once the OD_680_ reached 0.4^13^ cells were transferred into three separate 1000 ml Erlenmeyer flasks containing 400 ml of liquid TAP media. Cells were allowed to grow in constant conditions for two days. Once the OD_680_ reached 0.4, cells were used for the experiments, where 3 different flasks were treated as 3 independent biological replicates.

### Optimization of UV dose for efficient RNA-protein interactome capture in *C. reinhardtii*

To capture the in vivo RNA–protein interactome, we employed the orthogonal organic phase separation (OOPS) approach^75,76^, with minor modifications as described below. As this is the first application of OOPS in an algal species, we first optimized the UV irradiation dose to achieve maximum crosslinking efficiency, using crosslinking yield of both RBP and PBR as our benchmark (Ref). For in vivo UV crosslinking, approximately 200 million *C. reinhardtii* cells were pelleted by centrifugation at 1,000 × *g* for 10 minutes at 4 °C and resuspended in 10 mL of fresh, cold TAP medium. The cells were then transferred into 90 mm Petri dishes, placed on ice, and irradiated using a UVP CL-1000 crosslinker (Analytik Jena) equipped with 254 nm UV light at five different doses: 0.65, 1.95, 3.25, 4.55, and 5.85 Jcm^-^². The irradiation was applied in increments of 0.65 Jcm^-^², with a one-minute interval between each dose. Non-irradiated cells served as negative controls. Immediately after UV crosslinking, five crosslinked (CL) and one non-crosslinked (NC) samples from each plate were collected in 15 mL Falcon tubes and centrifuged at 5,000 × *g* for 10 minutes at 4 °C. The supernatant was discarded, and the cell pellets were resuspended in lysis buffer containing 10 mM Tris-HCl (pH 7.4), 500 mM NaCl, 1 mM EDTA (pH 8), 5 mM DTT, RNaseOUT (80 U; Invitrogen, Cat. No. 10777-019), and a complete protease inhibitor cocktail (Sigma, Cat. No. P9599). After thorough resuspension, cells were lysed at 19 kPa using a Constant Cell Disruption One-Shot (OS) model system. Lysates corresponding to 40 million cells in 100 µL from both CL and NC samples were mixed with 1 mL of PureZOL RNA isolation reagent (Bio-Rad, Cat. No. 7326890), vortexed for 15 seconds, and incubated at room temperature for 5 minutes. For phase separation, 200 µL of chloroform was added, followed by vigorous mixing for 15 seconds and a 5-minute incubation at room temperature. Samples were then centrifuged at 15,000 × *g* for 10 minutes at 4 °C. The upper aqueous phase, containing non-crosslinked RNA (free RNA), was carefully transferred to a fresh tube and processed for RNA isolation according to the manufacturer’s instructions. The interphase, containing crosslinked *C. reinhardtii* RNA-binding proteins (creRBPs) and protein-bound RNA (PBR), underwent two additional rounds of PureZOL purification. Finally, the interphase was precipitated using nine volumes of methanol. For digestion of protein-bound RNA (PBR), the interphase pellet was fully resuspended in 100 µL of 100 mM triethylammonium bicarbonate (TEAB) buffer (pH 8.5; Sigma-Aldrich, Cat. No. T7408). Samples were then incubated at 95 °C for 10 minutes with intermittent pipetting to ensure complete dissolution of the pellet. Once fully dissolved, CL and NC samples were incubated at 400 rpm for 16 hours with 4 µg of RNase A (Thermo Fisher Scientific, Cat. No. EN0531) and 10 U of RNase T1 (Thermo Fisher Scientific, Cat. No. EN0541). Subsequently, 1 mL of PureZOL reagent was added to each sample, followed by vigorous vortexing for 15 seconds. Phase separation was then performed using chloroform, as previously described. Following PBR digestion, creRBPs were released and partitioned into the lower organic phase. Approximately 450 µL of the organic phase was collected and subjected to precipitation using nine volumes of ethanol. The resulting protein pellet was resuspended in 50 µL of TEAB buffer (pH 8.5). creRBP concentrations were measured using Bradford assay (Bio-Rad, Cat. No. LIT33) with 10 µL of sample per reaction. We further assessed the impact of different UV doses on creRBP recovery via 12% SDS-PAGE followed by silver staining, allowing us to determine the optimal UV dose for crosslinking. These results were validated by immunoblotting with an antibody against a known creRBP, RBP40. To confirm the RNA-binding specificity of the recovered proteins, we again performed the RNase treatment and evaluated the results. For the RNase digestion assay, we employed the same UV dose as determined previously, with non-crosslinked (NC) samples serving as negative controls. Following UV crosslinking, 100 µL and 50 µL of crude lysate (corresponding to ∼40 million cells) were treated with RNase T1 and A in two separate tubes. After enzymatic digestion, creRBPs were recovered as described in a previous study^76^. The purified creRBPs were resolved on 12% SDS-PAGE gels, and subsequent silver staining revealed distinct protein profiles in RNase-treated and untreated samples. Additionally, we validated the purification strategy by analyzing the organic phases used during creRBP enrichment. Proteins were extracted from the first, second, and third organic phases, separated by 12% SDS-PAGE, and visualized via silver staining. We observed minimal to no protein signal in the third organic phase, indicating that proteins recovered in the final (fourth) organic phase following RNase treatment were not due to spillover or contamination from previous phases (Supplementary Figure S1A). This observation was further supported by immunoblotting with the RBP40 antibody, which showed no detectable signal in the third organic phase from either CL or NC samples (Supplementary Figure S1B), affirming the successful and specific capture of RBPs. To ensure that DNA-binding proteins were not co-purified, we probed for the well-characterized DNA-binding protein Histone H3 in both CL and NC samples across all four time points. As expected, H3 was not detected in any of the samples (Supplementary Figure S1C), further confirming the specificity of the creRBP purification protocol.

For the isolation of protein-bound RNA (PBR), the precipitated interphase from both crosslinked (CL) and non-crosslinked (NC) samples was incubated in 300 µL of Proteinase K (PK) buffer (10 mM Tris-HCl, pH 7.5; 1 mM EDTA, pH 8.0), supplemented with 12 units of Proteinase K (Merck, Cat. No. 1.24568.0100). The incubation was carried out for 2 hours at 50 °C to ensure complete protein digestion. Following digestion, PBR was purified using the phenol:chloroform:isoamyl alcohol extraction method (Sigma, Cat. No. 77617) according to the protocol described previously (Ref). The quantities of both free RNA and PBR were measured using a NanoDrop spectrophotometer (Thermo Fisher Scientific). RNA quality was assessed via electrophoresis on a 1% agarose gel, followed by UV imaging to evaluate rRNA integrity. Crosslinking efficiency was calculated using the captured total RNA and PBR with a formula that has been previously published (Ref). All procedures were performed in triplicate to ensure statistical reliability. To evaluate the recovery of free and protein-bound RNA at four different time points, two biological replicates were analyzed per time point. RNA recovery percentages were calculated using the protocol described above, and statistical significance was determined using a two-sided *t*-test comparing the total RNA yield between CL and NC samples (n = 8), as well as the percentage recovery.

### Circadian RBP and PBR Purification from *C. reinhardtii*

To synchronize the circadian clock, *Chlamydomonas reinhardtii* cells were cultured under a 12-hour light/12-hour dark (LD 12:12) cycle for a minimum of five days, as previously described. Following synchronization, the cells were transferred to constant light (LL) conditions at an intensity of 74 µmol m⁻² s⁻¹ for two days to allow progression under their free-running circadian rhythm. Three independently grown cultures were used as biological replicates. Cells were harvested in triplicate from each culture at four circadian time points: LL48, LL54, LL60, and LL66. Immediately after collection, the samples were UV-crosslinked at a dose of 4.55 J/cm². Non-crosslinked (NC) samples were collected in parallel at each time point to serve as negative controls. From both CL and NC samples at each time point, creRBPs were captured using the methodology described above, resulting in a total of 12 CL and 12 NC samples across the time course. The creRBP were quantified using Bradford (Bio-Rad, Cat. No. LIT33) and used for further downstream processing.

Similarly, protein-bound RNAs (PBRs) were isolated from three independent CL replicates, each representing one subjective day and one subjective night time point (LL48 and LL66). Due to the negligible yield of PBRs in NC samples, only crosslinked samples (total n = 6) were processed for downstream PBR analysis. The total RNA was quantified using nanodrop and sent for mRNA enrichment and illumina based-RNA-seq.

### SDS-PAGE, Western Blotting, and Silver Staining

An equal volume of RBPs isolated from non-crosslinked and crosslinked samples from 4-time points were boiled in a 1X Laemmli buffer at 95 °C for 10 minutes. After the incubation, proteins were separated on 12% SDS-PAGE using a mini protein gel running apparatus (BIO-RAD). RBPs in the gel were then visualized with silver staining^95^ or used for immunoblotting. Briefly, RBP proteins were transferred on the PVDF membrane using the BIO-RAD semi-dry western transfer apparatus at 10 mA for 7 minutes. Furthermore, the membrane was blocked for 1 hour at room temperature in 3 % nonfat dry milk (BIO-RAD) in 0.1% TBST buffer. For western blotting, RBP40 was used as a positive control. Anti-RBP40 (Agrisera Product no: AS14 2820) antibody was diluted in 3% non-fat dry milk at 1:1000 and incubated overnight at 4°C on a rocker with 50 rpm. After overnight incubation, the membrane was washed 5 times for 5 minutes each with 0.1% TBST followed by washing, the membrane was incubated for 1 hour at room temperature in goat anti-rabbit at 1:100,000 in the same blocking buffer (Thermo Fisher Scientific cat no-31460). Again, the membrane was washed 5 times with 0.1% TBST. Finally, the membrane was developed by using the ECL chemiluminescence substrate (BIO-RAD). Chemiluminescence blot images were taken in ChemDoc (BIO-RAD). The densitometry analysis was performed using ImageLab software (BIO-RAD) or in Fiji Image J. The dilution was 1:4000 for the AtpA (Agrisera,Product no: AS08 304) antibody, 1:1000 for the ADH (Agrisera, Product no: AS10 748) antibody, and 1:25,000 for the psbD antibody (Agrisera Product no: AS06 146). The western blotting was performed as per the manufacturer’s protocol. For the sliver staining of the immunoblots, we used a previously published protocol^96^.

### Proteomic sample preparation

creRBP pellets for each of the 24 samples were dissolved separately, each in 50 µl 0.1 M TEAB buffer, pH 8.5. Protein concentration was estimated using the Bradford assay (BIO-RAD-Cat. No. LIT33) and processed as described below.

#### a) Reduction, Alkylation, and Trypsin Digestion

Approximately 5 µg of protein from each sample was reduced with 10 mM dithiothreitol (DTT) at 56 °C for 45 minutes, followed by alkylation with 20 mM iodoacetamide (IAA) at room temperature for 45 minutes in the dark, as described previously (Ref). Proteins were then digested with sequencing-grade trypsin (Promega) at an enzyme-to-substrate ratio of 1:20 (trypsin:protein) for 16 hours at 37 °C with gentle agitation in a Thermomixer (Eppendorf) at 400 rpm. The digestion reaction was quenched by the addition of 0.5% trifluoroacetic acid (TFA). Peptides were desalted using C18 spin columns (Pierce) according to the manufacturer’s instructions. The desalted tryptic peptides were then vacuum-dried using a vacuum concentrator (Eppendorf) and reconstituted in 50% acetonitrile for downstream analysis.

### Spectral ion library generation

Tryptic peptides derived from the pooled protein samples were fractionated into eight fractions using a strong cation exchange (SCX) cartridge (Part No. 4326695) with a stepwise gradient of ammonium formate buffer (35–350 mM ammonium formate, 30% v/v acetonitrile, and 0.1% formic acid; pH 2.9). Peptides from each fraction were subsequently desalted using C18 ZipTips (Millipore, USA). Each fraction was then subjected to LC-MS/MS analysis using a quadrupole time-of-flight (QTOF) mass spectrometer (TripleTOF 6600, SCIEX) coupled to an Eksigent NanoLC-425 system. Optimized source parameters were applied: curtain gas and nebulizer gas were set to 25 psi and 20 psi, respectively; ion spray voltage was 5.5 kV; and source temperature was maintained at 250 °C. Approximately 4 µg of peptides were loaded onto a trap column (ChromXP C18CL, 5 µm, 120 Å, Eksigent, SCIEX) for online desalting at a flow rate of 10 µL/min for 10 minutes. Peptide separation was achieved on a reverse-phase C18 analytical column (ChromXP C18, 3 µm, 120 Å, Eksigent, SCIEX) using a 57-minute gradient at a flow rate of 5 µL/min. Mobile phases consisted of water with 0.1% formic acid (buffer A) and acetonitrile with 0.1% formic acid (buffer B). The gradient profile was as follows:

| **Time (min)** | **% A** | **% B** |
| --- | --- | --- |
| **0** | 97 | 3 |
| **38** | 75 | 25 |
| **43** | 68 | 32 |
| **45** | 20 | 80 |
| **45.5** | 10 | 90 |
| **48** | 10 | 90 |
| **49** | 97 | 3 |
| **57** | 97 | 3 |

Data was acquired using Analyst TF 1.7.1 Software (SCIEX). A 1.3-sec instrument cycle was repeated in high sensitivity mode throughout the entire gradient, consisting of a full scan MS spectrum (400–1250 m/z) with an accumulated time of 0.25s, followed by 20 MS/MS experiments (100–1500 m/z) with 50 msec accumulation time each, on MS precursors with charge state 2+ to 5+ exceeding a 120 cps threshold. The rolling collision energy was used, and the former target ions were excluded for 15 seconds.

### SWATH-MS data acquisition

The in-solution digested samples from the two groups were analyzed in SWATH-MS mode on the same instrument with similar LC gradient and source parameters as DDA runs. A SWATH-MS method was created with 100 precursor isolation windows, defined based on precursor m/z frequencies in a DDA run using the SWATH Variable Window Calculator (SCIEX), with a minimum window of 5 m/z. Accumulation time was set to 250 msec for the MS scan (400–1250 m/z) and 25 msec for the MS/MS scans (100–1500 m/z). Rolling collision energies were applied for each window based on the m/z range of each SWATH and a charge 2+ ion, with a collision energy spread of 5. Total cycle time was 2.8 sec

### Data analysis

A merged database search for DDA runs was performed using Proteinpilot^TM^ Software 5.0.1 (SCIEX) against *Chlamydomonas reinhardtii* proteome from UniprotKB (UP000006906, with 18,829 protein entries). Paragon algorithm was used to get protein group identities. The search parameters were set as follows: sample type-identification, cysteine alkylation-iodoacetamide, and digestion-trypsin. The biological modification was enabled in ID focus. The search effort was set to ‘Thorough ID’ and the detected protein threshold [Unused ProtScore (Conf)] was set to >0.05 (10.0%). False discovery rate (FDR) analysis was enabled. Only proteins identified with 1% global FDR were considered true identification. The group search result file from Proteinpilot^TM^ Software was used as a spectral ion library for SWATH analysis.

SWATH peak areas were extracted using SWATH 2.0 microapp in PeakView 2.2 software (SCIEX), and shared peptides were excluded. SWATH run files were added and retention time calibration was performed using peptides from abundant proteins. The processing settings for peak extraction were a maximum of 10 peptides per protein, 5 transitions per peptide, >95% peptide confidence threshold, and 1% peptide FDR. XIC extraction window was set to 5 min with 50 ppm XIC Width. All information was exported in the form of MarkerView (.mrkvw) files. In MarkerView 1.2.1 (SCIEX), protein area data was normalized, the data normalization strategy used was total area sum normalization and further analysis was performed using Python, R, MaxQuant Perseus, and Microsoft Excel.

### Proteomic Data Normalization and Statistical Data Analysis

Missing values were imputed using the MinDet function in the imputeLCMD package, with a threshold of q(MinDet) set to 0.01. To identify RNA-binding proteins (RBPs), we performed a two-sided t-test using the total area sum normalized data from three biological replicates at each of the four time points (LL48, LL54, LL60, and LL66). The analysis was conducted using MaxQuant Perseus software (version 1.6.14.0) with parameters set as follows: false discovery rate (FDR) of 0.05, s0 = 0.1, and 250 randomizations. Proteins with significant t-test results (FDR < 0.05 and log2 FC CL/NC ≥ 2 were classified as “creRBPs.” Additionally, proteins with 1≤ log2 FC CL/NC ≤ 2 and FDR < 0.05 from each of the four time points (CL/NC_LL_48, CL/NC_LL_54, CL/NC_LL_60, and CL/NC_LL_66) were designated as “candidate creRBPs”. A total of 518 creRBPs were identified and enriched in this process and the intensity of these 518 proteins exhibited a normal distribution pattern. Subsequent principal component analysis (PCA), Pearson correlation analysis, hierarchical clustering, and volcano plots were generated using either Python or the respective functions embedded within the Perseus software.

### Identification of temporal RNA binding proteins

To investigate the temporal dynamics of RNA binding among the creRBPs identified in the RNA-interactome, the total sum area normalized relative protein abundance was modeled using a cosine function (implemented in Perseus software) with a fixed period of 23.6 hours^17,97^. This free running period was selected based on previous studies. The amplitude and phase were treated as free parameters in the model. Using these parameters, we identified 372 creRBPs that displayed significant temporal changes in their RNA binding activity, with statistical significance indicated by a q-value < 0.05. Based on the normalized intensity values of these 372 dynamic creRBPs, hierarchical clustering was performed using the clustering package embedded in the SciPy library python (parameters: metric=’euclidean’, method=’average’,row_cluster=True, col_cluster=False), revealing two distinct groups: 200 creRBPs exhibited relatively high RNA-binding during the subjective day, while 172 creRBPs preferentially bound RNA during the subjective night. These temporal clusters were visualized using density plots, generated with the Seaborn package in Python.

### Identification of the dynamic binders

To identify “circadian dynamic binders,” we focused on 396 creRBPs that were common to both the circadian proteome^17^ and the current RNA-interactome study. We defined “circadian dynamic binders” as proteins whose temporal RNA-binding patterns are independent of their temporal abundance in the total proteome. To identify them we first calculated the differences between subjective night and subjective day time points (LL60 vs LL48, LL60 vs LL54, LL66 vs LL48, and LL66 vs LL54) in the RNA-interactome using a two-sided t-test with the following parameters: FDR ≤ 0.05, s0 = 0.1, and 250 randomizations. creRBPs that showed a fold change greater than 0.6 and an adjusted p-value ≥ 1 were considered to exhibit RNA-binding in a circadian pattern. To determine whether this circadian RNA binding was driven by changes in protein abundance, we performed the same pairwise comparisons using the total proteome data. The same parameters were applied to calculate differences between the corresponding time points in the total proteome. Using the python seaborn scatterplot function, we plotted the fold changes in the RNA-interactome and the total proteome with the total proteome time points on the x-axis and the RNA-interactome time points on the y-axis. This visualization allowed us to identify the “circadian dynamic binders.” Given that pairwise comparisons may result in overlapping protein IDs across subsets, we used UpSet plot function (embedded in the UpSetPlot library in python) to visualize and identify both common and unique proteins across different pairwise comparisons.

### Temporal Protein Bound RNA (PBR) Purification from *C. reinhardii*

The isolated PBR from the subjective day and subjective night time points were sent to Molsys Scientific Pvt. Ltd. for mRNA purification followed by mRNA - seq. using illumina Novoseq 6000 platform.

### mRNA Sequencing, RNA-Seq data processing, and Bioinformatics

Quality assessment of the raw FASTQ reads was performed using FastQC v.0.11.9 (default parameters). The raw FASTQ reads were pre-processed with Fastp v.0.20.1, followed by quality reassessment using FastQC and summarization via MultiQC. The indexed genome of *Chlamydomonas reinhardtii (Chlamydomonas reinhardtii strain CC-503 cw92 (NCBI ID: NC_057004.1))* was then mapped to the processed reads using HISAT2. Aligned reads were converted to BAM format and sorted using Samtools v.1.7. The aligned reads from individual samples (three subjective day-LL48 and three subjective night-LL66) were quantified using featureCounts v.0.46.1 (parameters: -g gene_id -F GTF -p for gene counts and -g transcript_id -F GTF -p for transcript counts). These gene and transcript counts were used as inputs for DESeq2 to estimate FPKM (Fragments per Kilobase of transcript per Million mapped reads) values. The FPKM values were exported in Excel format for further downstream analysis.

In total, 17,742 transcripts were identified. The FPKM values were log-transformed, and missing values were imputed using the MinDet function (q = 0.01) in MaxQuant Perseus software. Data were filtered to retain valid values greater than 0 in at least 70% of the replicates, resulting in 2,476 transcripts. The FPKM values were then quantile-normalized using MaxQuant Perseus software. Subsequent analyses, including principal component analysis (PCA), volcano plots, hierarchical clustering, and other statistical assessments, were performed using MaxQuant Perseus or Python.

#### 2) Gene ontology analysis

Gene ontology analysis of the RNA-binding proteins (RBPs) and the protein-bound RNAs (PBRs) was conducted using the PANTHER Classification System (version 17.0; http://www.pantherdb.org/)^98^. The analysis was performed against the *Chlamydomonas reinhardtii* reference proteome (release 2023_03), which comprises 17,613 protein entries. Functional enrichment was assessed using Fisher’s exact test, and significance was tested using the Benjamini–Hochberg false discovery rate (FDR) method. GO terms with an adjusted p-value (FDR) less than 0.05 were considered significantly enriched. Fold enrichment values and corrected p-values for each GO term were visualized using R or Python-based data analysis pipelines. To compare GO term coverage across species, datasets of RBPs identified in other organisms were obtained from RBPbase (version 0.2.1 alpha; https://apps.embl.de/rbpbase/). This comparative analysis allowed for cross-species functional assessment of RNA-binding proteins.

## Data availability

The mass spectrometry proteomics raw data have been deposited to the ProteomeXchange Consortium via the PRIDE^99^ partner repository with the dataset identifier PXD064344. Along with the raw data we have also uploaded the excel files with all details. The transcriptomics raw data have been deposited to NCBI SRA under the BioProject ID: PRJNA1269807 with the dataset identifier SUB15312724.

## Acknowledgements

The authors thank Dr. Shantanu Sengupta and Mr. Praveen Singh of the National Facility for Biochemical and Genomic Resources (**NFBGR**), IGIB for help with SWATH-MS sequencing.

## Funding

This study was supported by the Science and Engineering Research Board, Government of India (SERB grant SRG/2019/000364) and Department of Biotechnology, Government of India (DBT grant BT/PR32511/BRB/10/1803/2019), SR acknowledges the Annual Research Grant from Ashoka University. DBJ was supported by a PhD fellowship from Ashoka University.

## Author contributions

**Dinesh Balasaheb Jadhav:** Experimental designing; data curation; software; experimentation - designing and performing; formal analysis; validation; investigation; visualization; methodology; writing –review and editing. **Sougata Roy:** Conceiving the original idea; experiment designing; resources; supervision; project administration; formal analysis-some: funding acquisition; writing – original draft; writing – review and editing.

## Disclosure and competing interests’ statement

The authors declare that they have no conflict of interest.

## Notes

### Competing Interest Statement

The authors have declared no competing interest.

### Summary of Updates

Only the ORCID ID of the co-author has been added.

