## Supplementary Figures for "Temporal dynamics of RNA metabolic enzyme interactome lay the foundation for riboregulation mediated circadian metabolism"

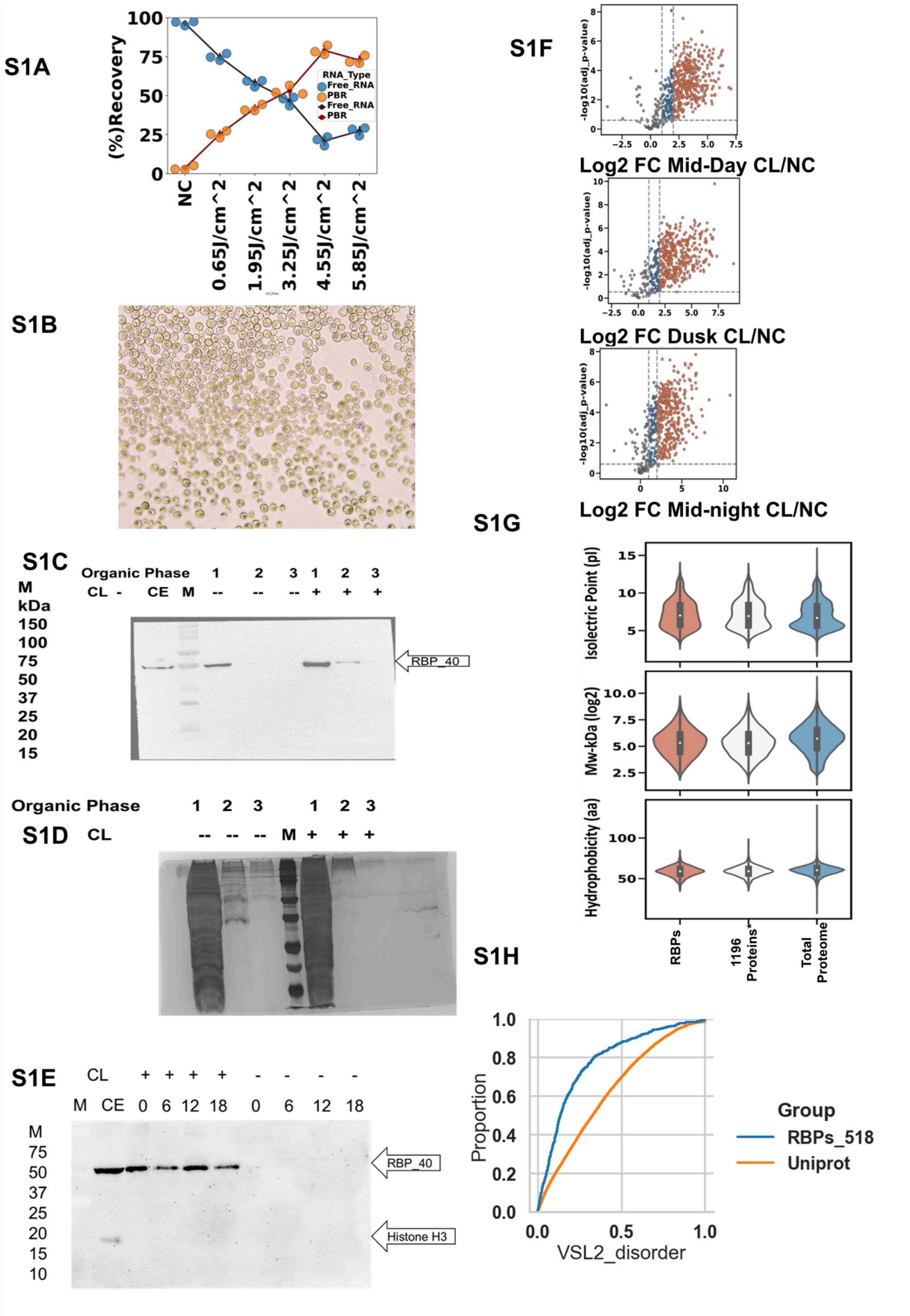


**Supplementary Figure 1: Optimization of the RIC in *C. reinhardtii.* S1A)**

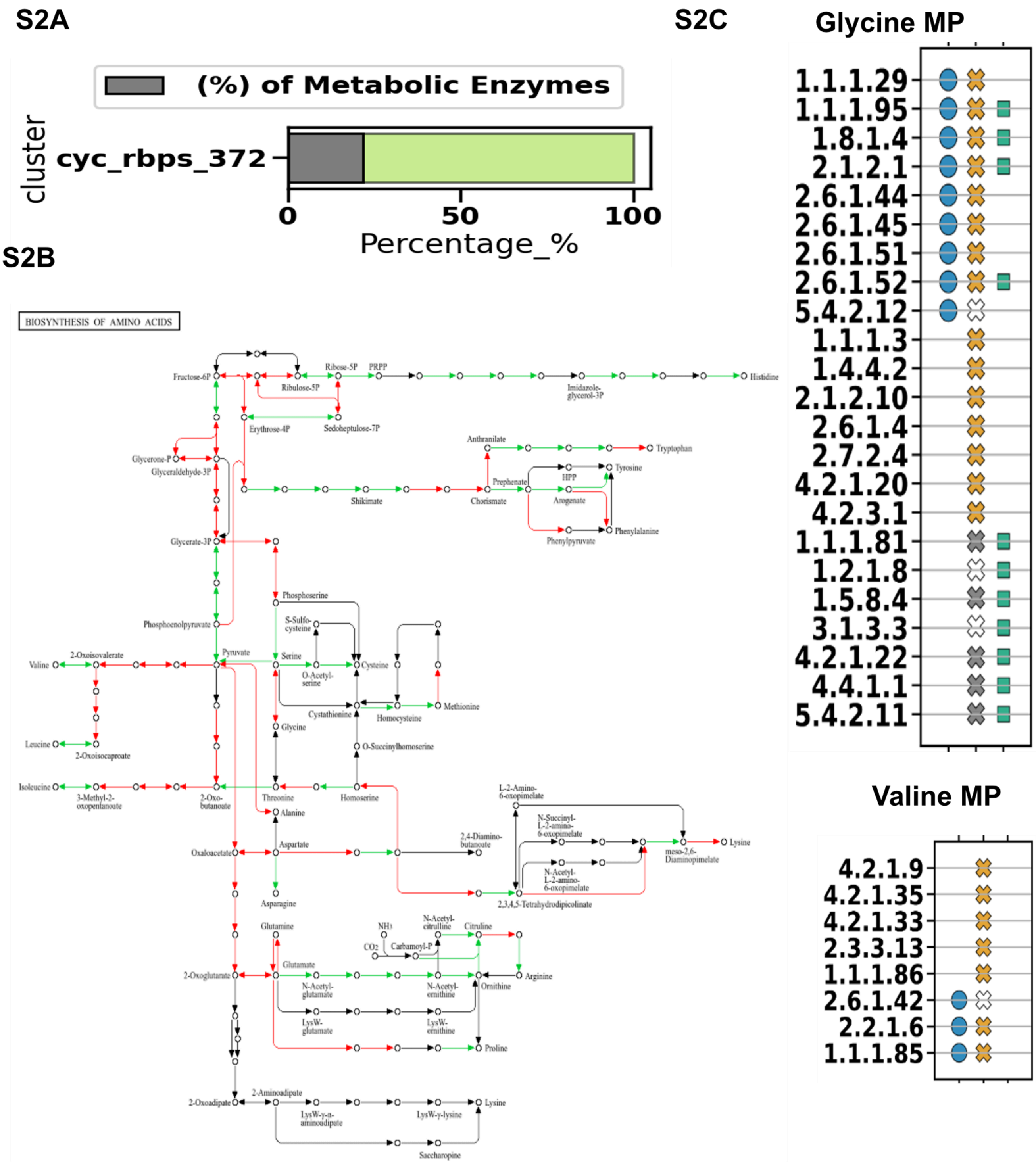


**Supplementary Figure 2: Enrichment of metabolic enzymes from the Amino acid metabolic pathways.**

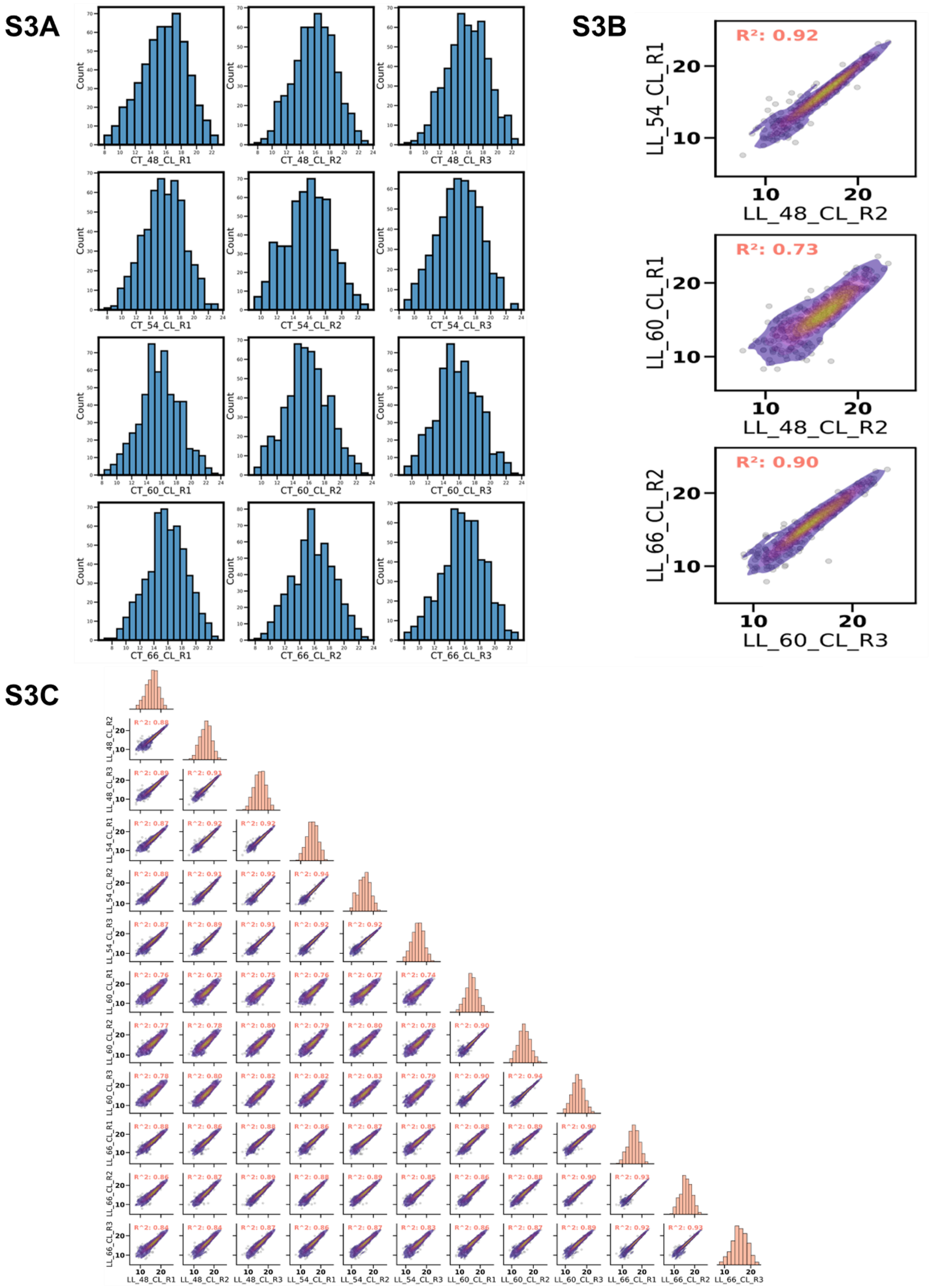


**Supplementary Figure 3: Data quality of the the creRBP protein enrichment after mass spectrometry.**

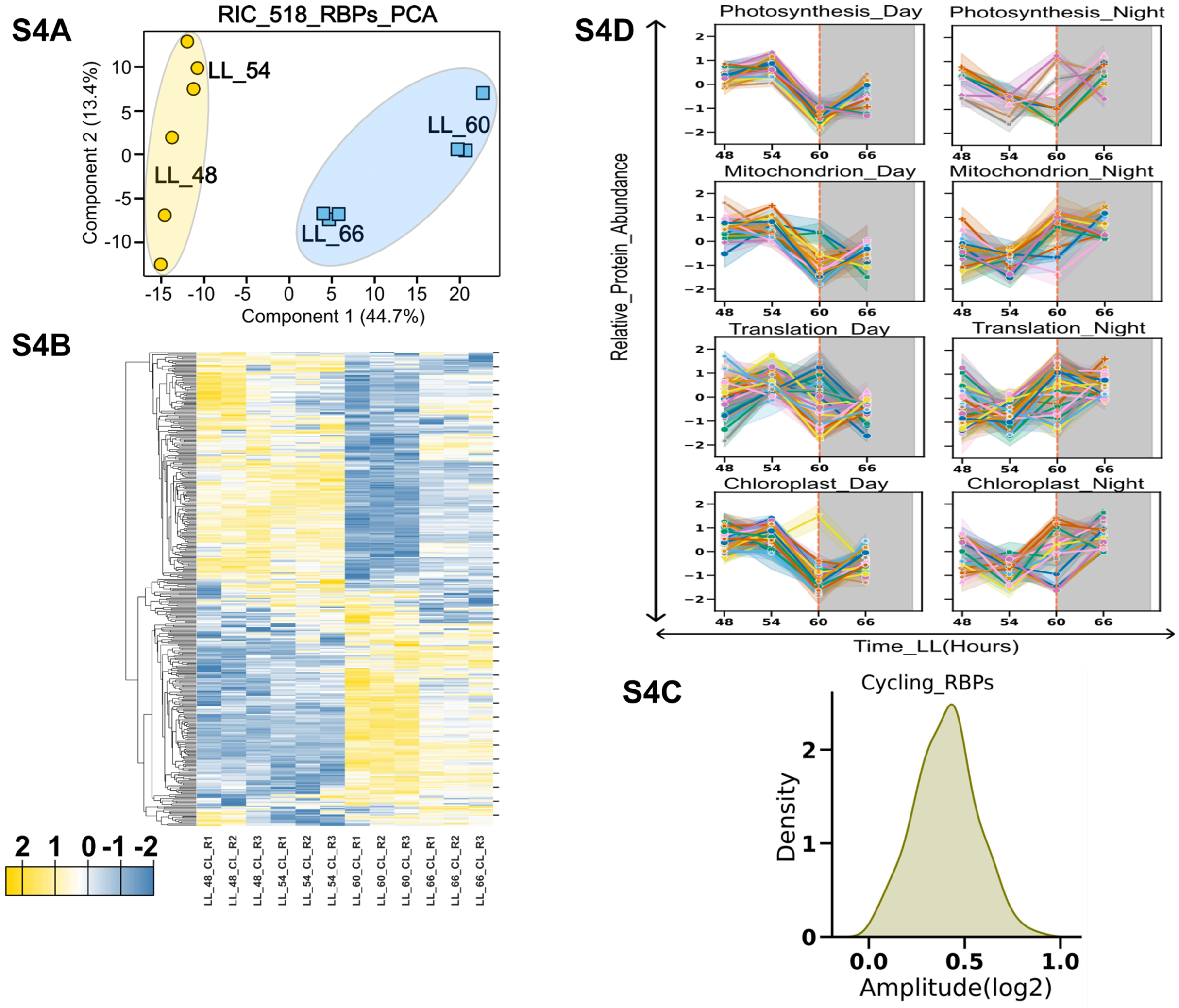


**Supplementary Figure 4: creRBP exhibiting circadian dynamicity in RNA binding.**

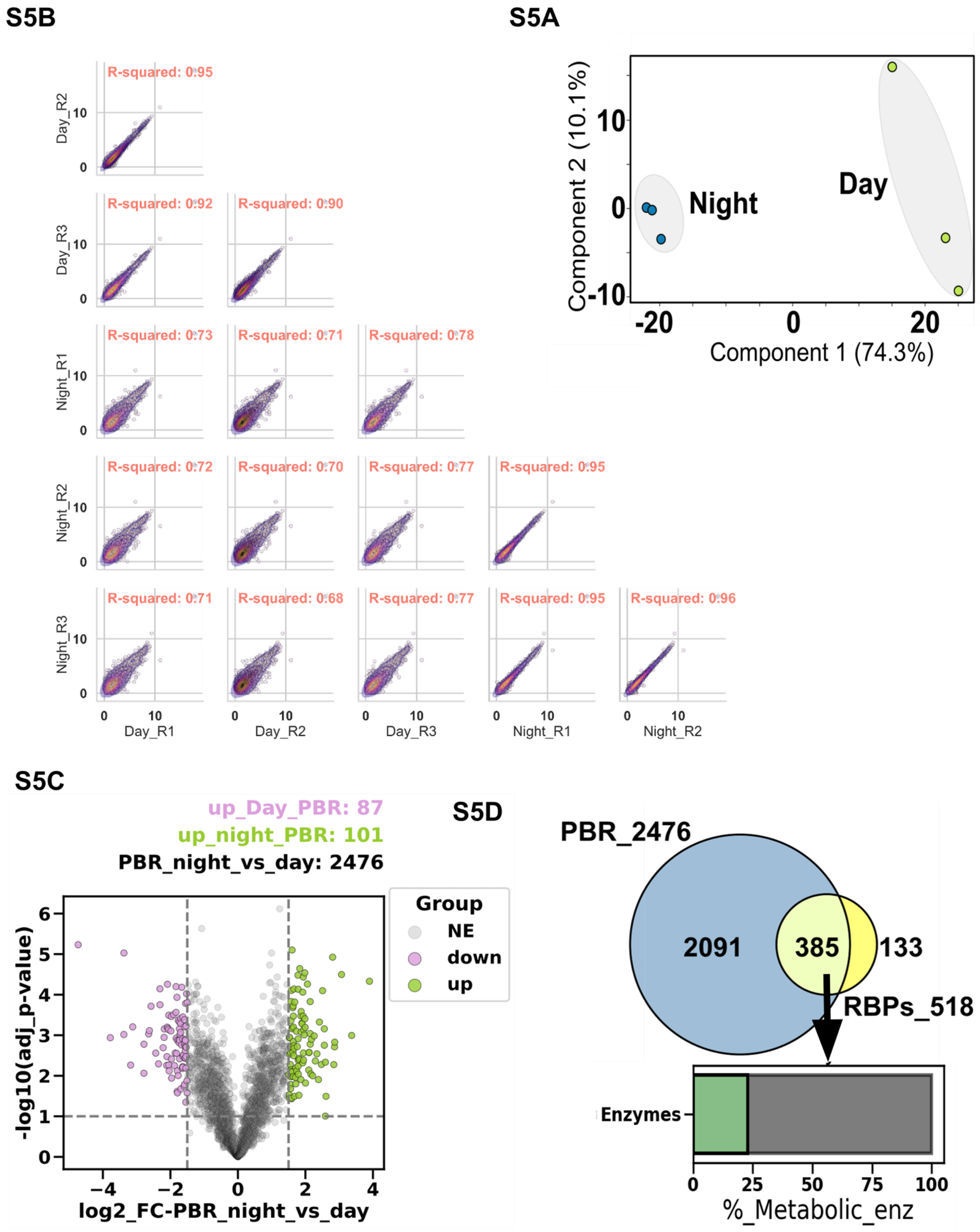


**Supplementary Figure 5: Temporal dynamics of the protein bound RNA (PBR) captured from *C. reinhardtii* grown under constant light.**

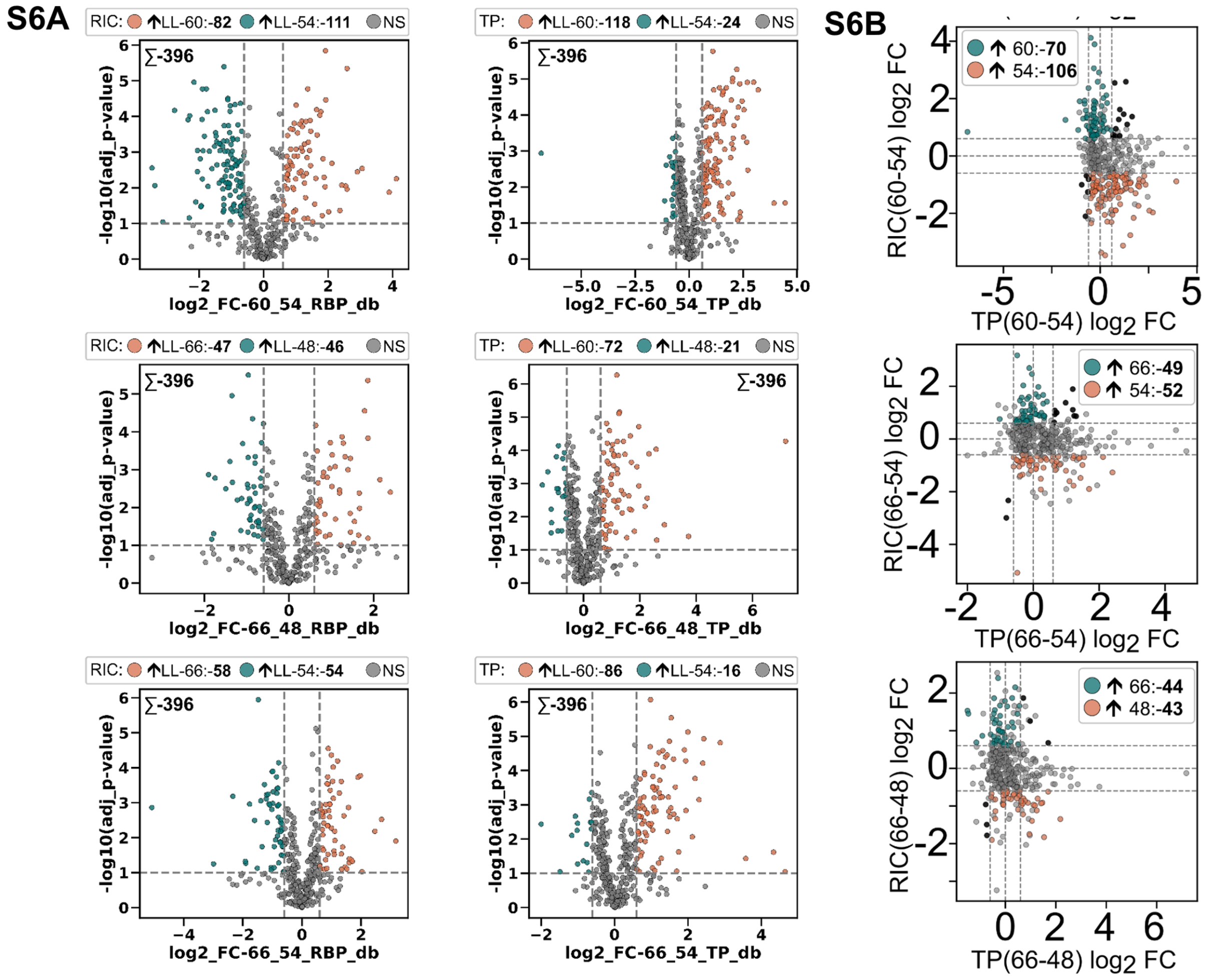


**Supplementary Figure 6: Circadian dynamic binders from subjective day and subjective night.**

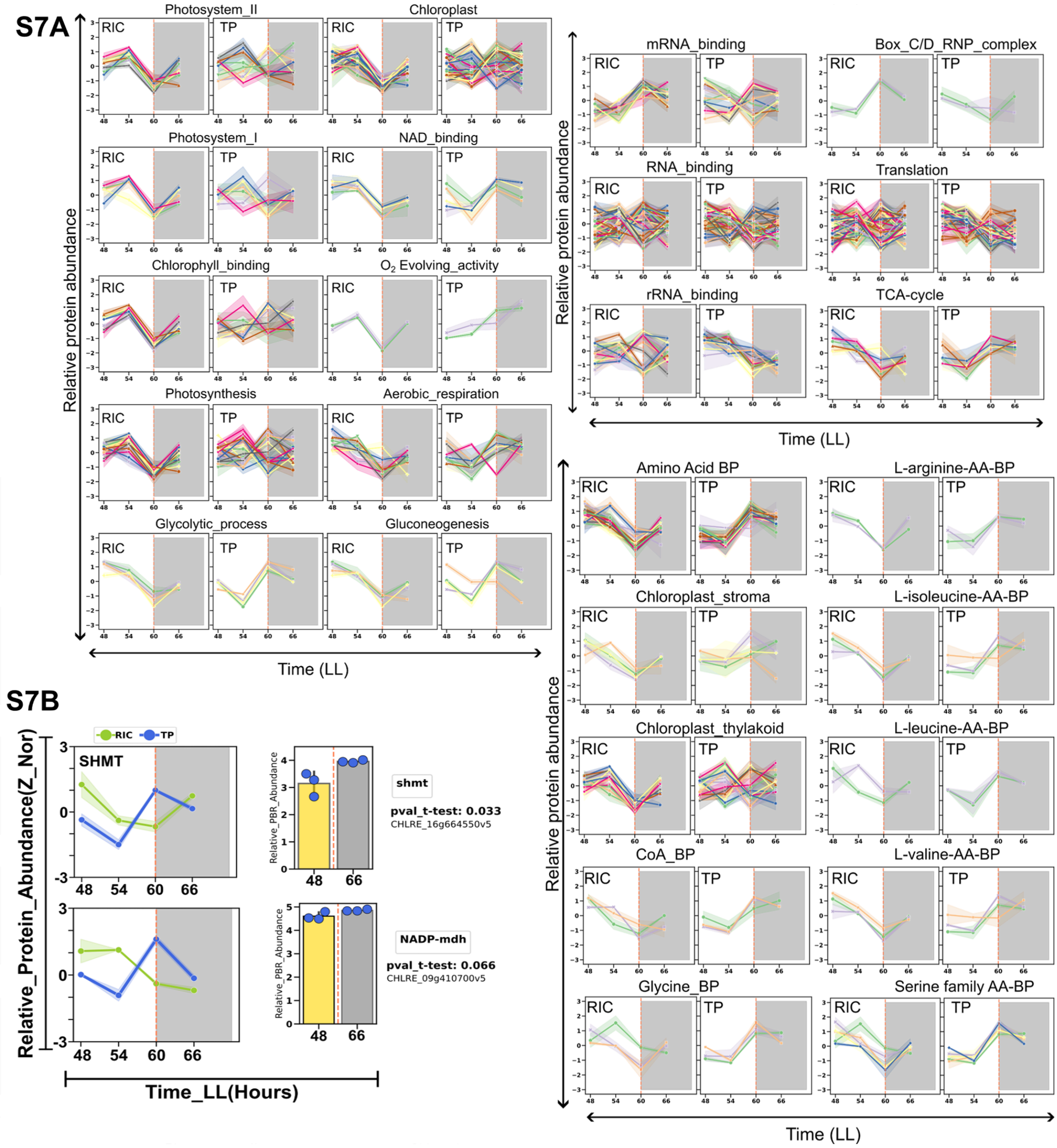


**Supplementary Figure 7: Metabolic enzymes and proteins from different cellular processes and organelles that are classified as circadian dynamic binders.** .
